# Intra-subtype heterogeneity shapes treatment response in *KMT2A*-rearranged ALL across all age groups

**DOI:** 10.1101/2025.06.05.657589

**Authors:** Alina M. Hartmann, Lorenz Bastian, Malwine J. Barz, Johannes Haas, Eric Amelunxen, Patrick Ehm, Lennart Lenk, Michaela Kotrova, Thomas Beder, Fabio Steffen, Kerstin Rauwolf, Nadine Wolgast, Sonja Bendig, Cecilia Bozzetti, Julia Alten, Mayukh Mondal, Annika Rademacher, Julia Heymann, Wencke Walter, Claudia Haferlach, Aeint-Steffen Ströh, Anke K. Bergmann, Thomas Burmeister, Nicola Gökbuget, Beat Bornhauser, Jean-Pierre Bourquin, Monika Brüggemann, Martin Schrappe, Gunnar Cario, Claudia D. Baldus

## Abstract

**Background:** *KMT2A*-rearranged B-cell acute lymphoblastic leukemia (*KMT2A*r B-ALL) exhibits significant heterogeneity in age of onset, developmental origins, and clinical outcomes. The interplay of individual factors influencing early treatment response within this high-risk molecular subtype remains poorly elucidated. We aimed to comprehensively assess how leukemic developmental state, fusion partner, and patient age jointly influence early treatment response and drug sensitivity.

**Methods:** To identify determinants of early treatment response to induction chemotherapy, we analysed 465 *KMT2A*r B-ALL cases spanning a wide age range (1 month to 89 years) by integrating transcriptomic and genomic profiling with functional drug response and measurable residual disease (MRD) kinetics. Transcriptomic profiling was used to derive a developmental maturity score based on proximity to physiological B-cell differentiation from a normal B-lymphopoiesis reference. Using an ordinal regression model we identify gene expression programs associated with early MRD response.

**Results:** We observed a strong inverse correlation between MRD clearance with advancing age (p=2.1E-04), proximity to early B-cell-precursor developmental state (low maturity score, p=1.3E-03) and *AFF1* as fusion partner (p=7.0E-04). A combined model confirmed the predominate impact of both maturity and *KMT2A* fusion partner on MRD response, supporting the concept that the cell’s developmental state defines therapy response. Gene expression analysis identified cellular traits that relate to MRD response (e.g. chromatin organization, immune modulation and proliferation). This gene expression classifier grouped cases by MRD response but also by *ex-vivo* induction drug sensitivity. Notably, good responders to *ex-vivo* induction drugs were characterized by a higher maturity score (p=1.8E-03), whereas for less mature *KMT2A*r B-ALL cases response profiles suggested higher Venetoclax sensitivity.

**Conclusions:** Our study provides an integrative framework linking developmental phenotype, fusion partner, and MRD kinetics across the full age spectrum of *KMT2A*r B-ALL. The maturity score, derived from bulk transcriptome data, offers a biologically relevant predictor of early treatment response and drug sensitivity. These insights may support future risk-adapted strategies and therapeutic targeting, particularly in immature *KMT2A*r B-ALL.

## Introduction

*KMT2A*-rearranged B-cell acute lymphoblastic leukemia (*KMT2A*r B-ALL) is considered a high-risk molecular subtype due to poor treatment response and high risk of relapse. *KMT2A* is the most prevalent genomic driver in infant ALL, accounting for ∼70% of the cases(1,2), yet also occurs in older children (∼2-5%(2,3)) and adult patients (∼15%(4)). The prognosis is especially poor for infant patients and older adults, suggesting intra-subtype molecular heterogeneity to account for the differences in treatment response. Over 90 *KMT2A* fusion partners have been identified with over 80% of *KMT2A*r cases having an *AFF1*, *MLLT1* or *MLLT3* rearrangement(5). Adult cases predominantly carry *KMT2A::AFF1* rearrangements (*AFF1*r); pediatric and infant cases display a greater variety of fusion partners. *KMT2A* fusion partners have shown to impact therapy response and outcome with *MLLT3*r cases showing superior outcome compared to *AFF1*r in infant patients, but not in pediatric patients (age >1 year)(2,6). Molecular characterization of an infant *KMT2A*r B-ALL cohort revealed subclusters associated with different driver fusions(7), suggesting different mechanisms based on the underlying gene fusion partner.

In addition to the heterogeneity of molecular drivers, molecular B-ALL subtypes have differential proximity to the physiological developmental stages of normal B-cell development, from hematopoietic stem cells (HSCs) to committed pre-B cells. Age-overriding immunoglobulin rearrangement profiling across molecular subtypes revealed a higher immunogenetic maturity in pediatric patients, compared to adults(8,9) and single-cell-sequencing studies of ALL found increasing cell plasticity with age(10). However, molecular characterization of *KMT2A*r B-ALL cases have unravelled an immature leukemic origin in younger patients(10–12) associated with increased cell plasticity, retained latent myeloid potential and co-expression of myeloid marker genes(10). This immature, high plasticity phenotype has been shown to impact drug sensitivity(13), (14), necessitating alternative treatment strategies. As the bispecific T-cell engager monoclonal antibody, blinatumomab, chimeric antigen receptor T-cell therapy and novel agents such as BH3-mimetic Venetoclax and Menin inhibitors(15,16) are being incorporated into novel treatment algorithms, futures investigations will have to assess their efficacy in the light of the underlying leukemic origin of the ALL.

To address the multi-faceted intra-subtype heterogeneity in an age-overarching cohort and to explore the impact of the underlying cellular context in disease and therapy response, we conducted a multi-omics analysis in a large cohort of 465 *KMT2A*r diagnostic B-ALL cases spanning a unique age range upon diagnosis (1 month to 89 years), and including common *KMT2A* fusions with representative frequencies(5). We demonstrate that the maturity score (reflecting proximity to B-cell developmental stages), the *KMT2A* fusion partner and the age at diagnosis impact measurable residual disease (MRD) response in *KMT2A*r ALL. A machine learning model revealed maturity and fusion-partners as stronger predictors for MRD than age. Importantly, distinct leukemic regulatory programs are linked to treatment response and thus may provide the basis for understanding and targeting molecular mechanisms of drug resistance in *KMT2A*r B-ALL.

## Methods

### Patients and transcriptome data set

We have aggregated a unique cohort of n=465 *KMT2A*r ALL cases spanning the whole age range (1 month to 89 years, Figure 1A) including most common *KMT2A* fusion partners with representative frequencies(5) (Figure 1B). The cohort is well characterized with available omics layers including RNAseq (n=325), Sanger sequencing/RNAseq for gene fusion detection (n=436), SNP arrays (n=135), lymphoid capture panel DNA-sequencing(17) (n=82) and drug response profiling(18,19) (DRP, n=61). Moreover, data on clinical annotations including age (n=422), genomic *IG-R* rearrangement profile (n=47), white blood cell count at diagnosis (WBC, n=223) and longitudinal MRD monitoring (n=214) were available. For the n=325 RNAseq cases, n=148 were uniformly sequenced in our facility and defined as discovery cohort. The remaining n=177 cases (validation cohort) were from publicly available sources(20) (n=133) and from a collaborating laboratory(21) (n=41). (Figure 1A; Supplementary Table S1)

**Fig. 1.**
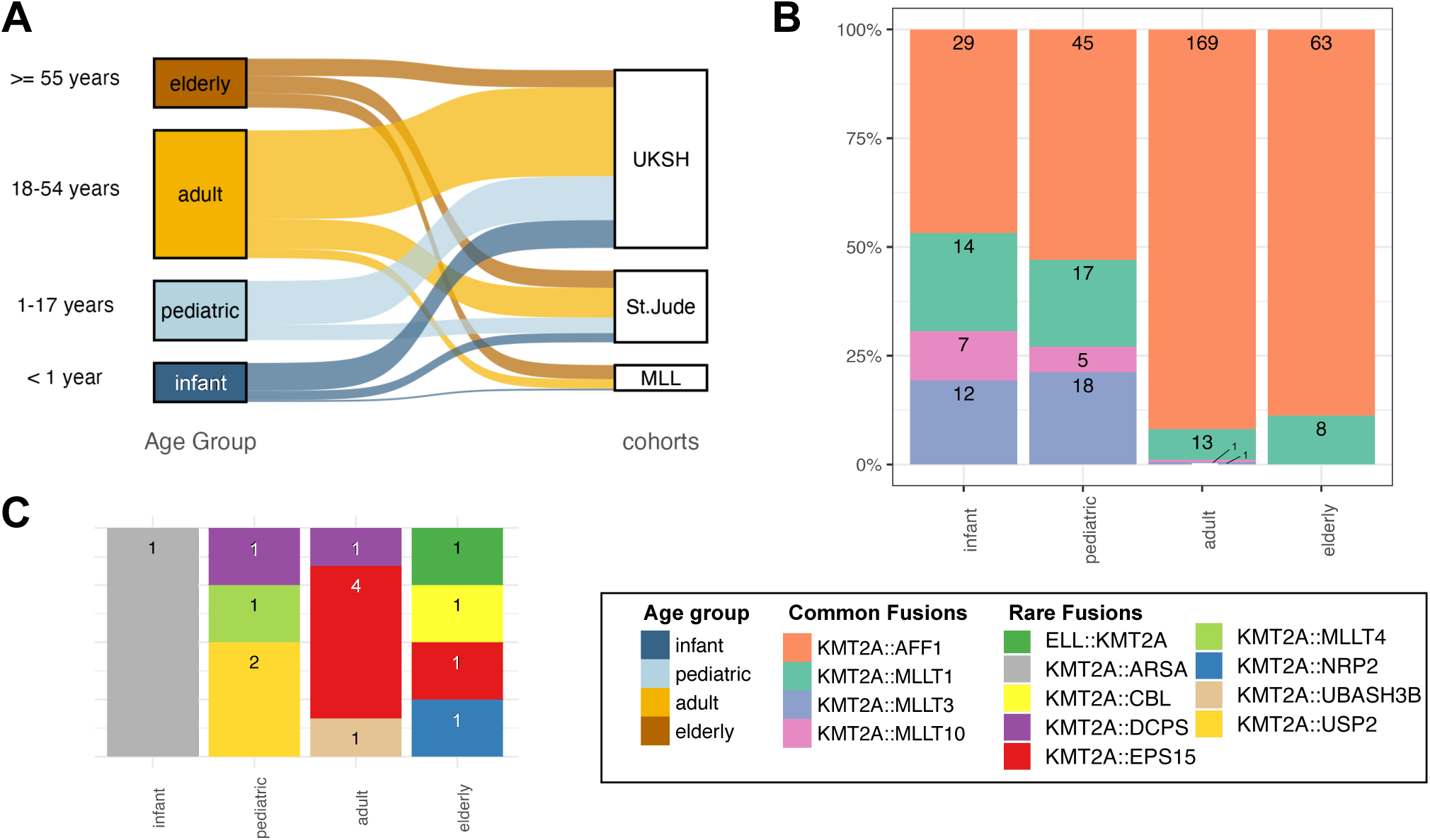
Age-overriding *KMT2A*r B-ALL patient cohort. **A**: Included patients (n=465) of different age groups (>=55 n=79, 18-54 n=207, 1.17 n=96, <1 n=63) from three cohorts (UKSH n=288, St. Jude(20) n=136, MLL(21) n=41). **B:** Distribution of common KMT2A fusions across age groups (n=422, 96.6%). **C**: Distribution of rare KMT2A fusions (n=15, 3.4%).

### Multi-omics data

Transcriptomic data analysis, fusion gene detection and DNA capture panel sequencing were performed by common standard (Supplementary Methods). DRP was performed in an established co-culture system as previously described(18,19,22) (Supplementary Methods).

### Clinical response data

For patients treated according to the AIEOP-BFM ALL/INTERFANT and GMALL study group protocols, MRD measurements were performed in the central reference laboratories. Pediatric-inspired induction therapy protocols in adults with comparable MRD timepoints in pediatric regimens allowed for common kinetic definitions of MRD clearance (Supplementary Methods, Supplementary Figure S1).

### Data analysis and statistics

We identified *KMT2A* fusion specific gene sets with multi-comparison ANOVA and LASSO feature selection to define genes uniquely expressed in patients with *AFF1*r, *MLLT1*r or *MLLT3*r. The MRD gene expression signature was defined using an ordinal regression model to account for the structured nature of MRD categories (Supplementary Methods).

To validate gene sets, we performed unsupervised clustering of normalized gene counts across our discovery cohort and, where applicable, the validation cohort, with the R pheatmap package (Version 1.0.12). Biological function was annotated using GO-term enrichment analysis (Supplementary Methods).

To define the patient-specific maturity score, representing proximity to physiological B-cell developmental states, we used the machine learning classifier ALLCatchR(23) to define developmental trajectories and fitted linear slopes on the enrichment across B-cell developmental stages. High maturity score values indicate proximity to mature B-cell stages, low maturity scores correspond to more immature B-cell stages. To validate our maturity score we mapped our cohort to two recently published single-cell atlases of physiological B-cell development(10,24) and additionally correlated it to the B-cell map author’s multipotency score(10) (Supplementary Methods).

To quantify how age, maturity and gene fusion effect MRD response, we trained a machine learning decision tree (Supplementary methods). The model did not have high enough predictive capacity to be used as an MRD-clearance classification tool (combined accuracy 56.7%, compared to 27.9% random assignment). However, feature importance values could be extracted to learn which features contribute most to the decision process. We validated these findings in a univariate odds ratio analysis (Supplementary Methods).

### Data Availability

RNAseq count data are accessible from zenodo at doi 10.5281/zenodo.15638437 for our cohort or through the original publications (St.Jude(20) and MLL(21)). DRP normalized AUC data are provided in Supplementary Table S7. RNAseq raw fastq files will be provided upon acceptance.

## Results

To disentangle the molecular underpinnings of treatment response in *KMT2A*r B-ALL, we aggregated a cohort of n=465 initial diagnosis samples, including n=325 RNAseq samples from own (n=148) and external (n=177)(20,21) sequencing, representing three clinical cohorts and an age spectrum from 1 month to 89 years (Figure 1A). Patients were categorized for subsequent analyses as infants (<1 year; n=63), pediatric (1-17 years, n=96), adult (18-54 years, n=207) and elderly patients (>=55 years, n=79). To avoid batch effects, we processed n=148 samples homogeneously in-house (patient age: 2 months to 79 years, median 13 years), serving as a discovery cohort. Integrative data analysis comprised genomic profiling for karyotypes (SNParray), single nucleotide variants (capture panel sequencing(17)), and IG/TR rearrangement status, transcriptomic profiling for gene expression and gene fusions and functional DRP.

### *KMT2A* fusion partners are associated with age and characterized by few cooperating genomic aberrations

Among B-ALL molecular subtypes, *KMT2A*r ALL is characterized by the widest age range and holds a diverse repertoire of *KMT2A* fusion partner genes. We observed a significant correlation between fusion partners and patient’s age (Figure 1B; n=305, Chi-sq. test P<0.0001). While *AFF1* was the most frequent fusion partner gene (n=221/321; 69%), it was strongly enriched in adult/elderly patients (71% vs infant/pediatric 48%; p=7.7E-12). Pediatric patients showed a more diverse fusion partners, with *MLLT3*r exclusively found in infants/pediatric cases, and *MLLT1*r and *MLLT10*r enriched in infant/pediatric cases (61% and 92% vs adult/elderly 8% and 0.6%, p=0.008 and p=1.5E-4, respectively). Rare *KMT2A* fusion partners were also included in our cohort [*EPS15* (n=5), *USP2* (n=2), *DCPS* (n=2), one of each: *MLLT4*, *UBASH3B*, *NRP2*, *ARSA*, *CBL*, *ELL,* Figure 1C] providing a comprehensive representation of *KMT2A*r ALL.

To investigate the landscape of cooperating genomic events, we performed capture panel sequencing for genes recurrently involved in ALL (n=82) and SNParrays (n=104). We detected pathogenic mutations in 32 of the 82 analysed samples [in 8/26 (31%) pediatric patients and in 24/56 (43%) adult patients]. In total, 21 of 82 patients (26%) carried either a KRAS or NRAS mutation, the remaining mutations were found in TP53 (6%), ATM (5%), CDKN2B (2%) and CXCR4, FAT1, RUNX1, TCF3 (<2%, Supplementary Figure S2A). Mutations were neither enriched in age groups nor associated with driver fusions. The SNP arrays revealed diploid karyotypes in nearly all cases, without patterns of recurrent chromosomal gains and losses (Supplementary Figure S2B).

### Gene regulatory landscape of *KMT2A*r fusion partners

Multi-comparison ANOVA and LASSO feature selection were used to define genes uniquely expressed in *AFF1*-, *MLLT1*- or *MLLT3* fusion positive cases. We unravelled a 237-gene signature in our discovery cohort. The gene signature separated *AFF1*r cases from other *KMT2A* fusions in our discovery cohort (Figure 2A) and in the two independent validation cohorts(20,21) (Supplementary Figure S3A and S3B respectively). *MLLT1* and *MLLT3* fusions were separated only at the second clustering level, indicating that these cases are more similar to each other than to *AFF1*r cases (Figure 2A).

**Fig. 2.**
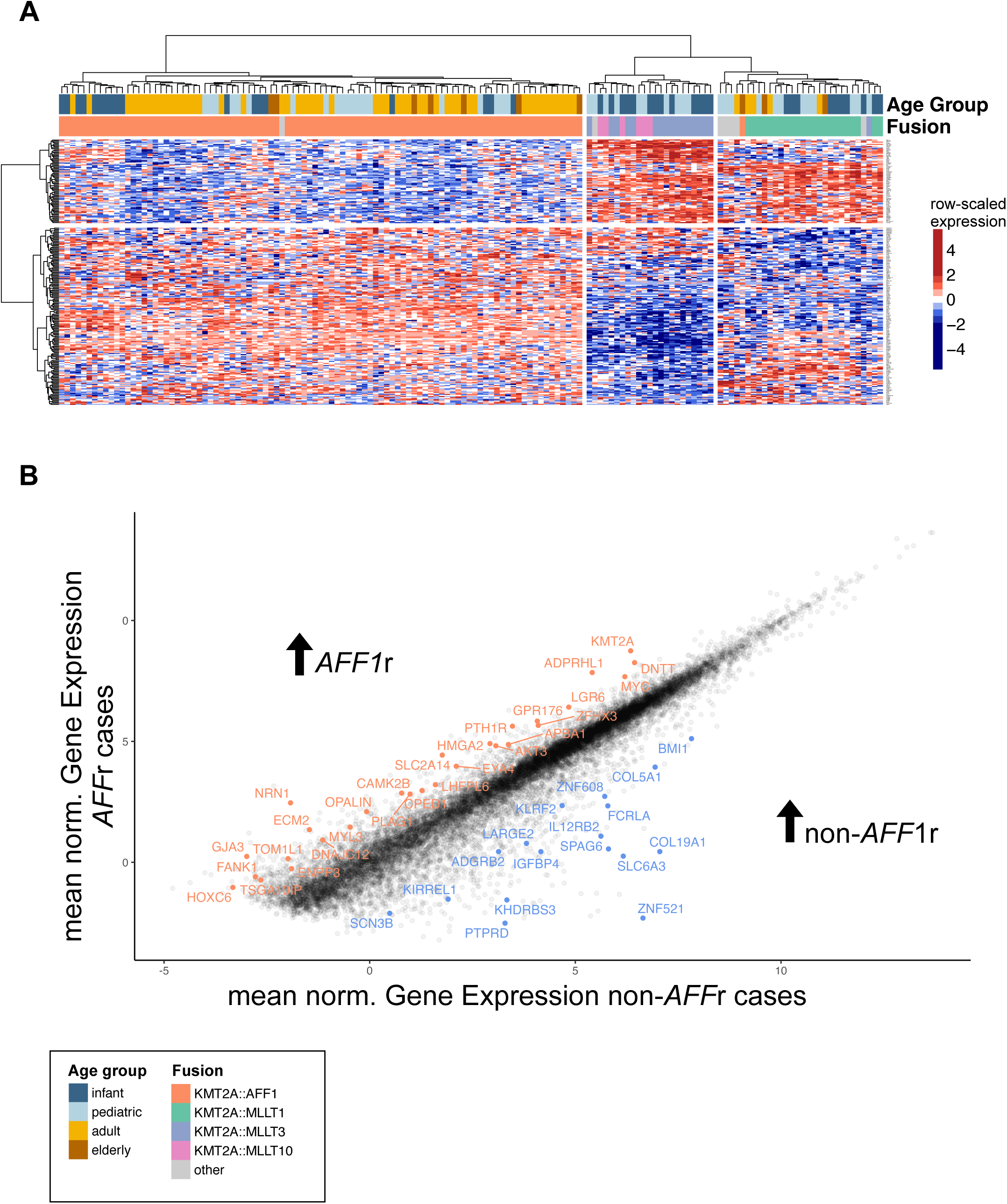
Gene regulatory landscape of *KMT2A*r fusion partners. **A**: Heatmap of hierarchical clustering of fusion specific genes detected in supervised analysis using multi-comparison ANOVA and subsequent LASSO feature selection for genes separating *AFF1*r, *MLLT1*r and *MLLT3*r (n=148 samples, n=236 genes). Hierarchical clustering splits *AFF1*r and non-*AFF1*r patients, the second split subdivided non-*AFF1*r cases **B**. Mean gene Expression (norm. log2CPMs) per gene of *AFF1*r (y-axis) vs. non-*AFF1*r (x-axis) cases. Highlighted Genes represent genes that were detected by ANOVA and LASSO feature selection (panel A). Genes significantly higher in *AFF1*r cases at log2fc >1.5 are highlighted in orange (above diagonal), genes significantly higher expressed in non-*AFF1*r cases at log2fc >2.5 are highlighted in blue (below diagonal).

Functionally, this gene signature differentiated *AFF1*r cases from non-*AFF1*r cases. Upregulated genes in *AFF1*r cases included leukemia-associated proliferation genes *MYC*, *AKT3* and *KMT2A* itself, other known cancer genes (e.g. *HMGA2*, HOXC6, *PLAG1*) and early stem-cell markers such as *DNTT* and *LRG6* (Figure 2B – above diagonal). This signature indicates the activation of highly proliferative, stem-cell-near cellular programs with globally activated gene expression and somatic recombination as well as silenced apoptosis and differentiation signals suggestive of an immature gene regulation in *AFF1*r cases.

In contrast, genes upregulated in non-*AFF1*r cases included B-cell functional genes like fc-like receptor protein *FCRLA, IGFBP4* and *ZNF521* and proliferation activation genes like *BMI1* and *IL12RB2,* revealing a more mature, differentiated and functionally active B-cell program that is nonetheless proliferating (Figure 2B – below diagonal).

### *KMT2A*r ALL in adults is enriched for immature transcriptional developmental states

To systematically characterize the developmental underpinnings of *KMT2A*r ALL, we made use of our immuno-genomic defined RNAseq reference of human B-lymphopoiesis(23). Single sample gene set enrichment analysis (ssGSEA) for gene sets defining normal B-lymphopoiesis was used to define the proximity of individual *KMT2A*r ALL samples to their physiological B-cell counterparts (Figure 3A). Regression analysis across enrichment scores was used to condense values into a single maturity score (Figure 3B, Supplementary Figure S4A, Material & Methods).

**Fig. 3.**
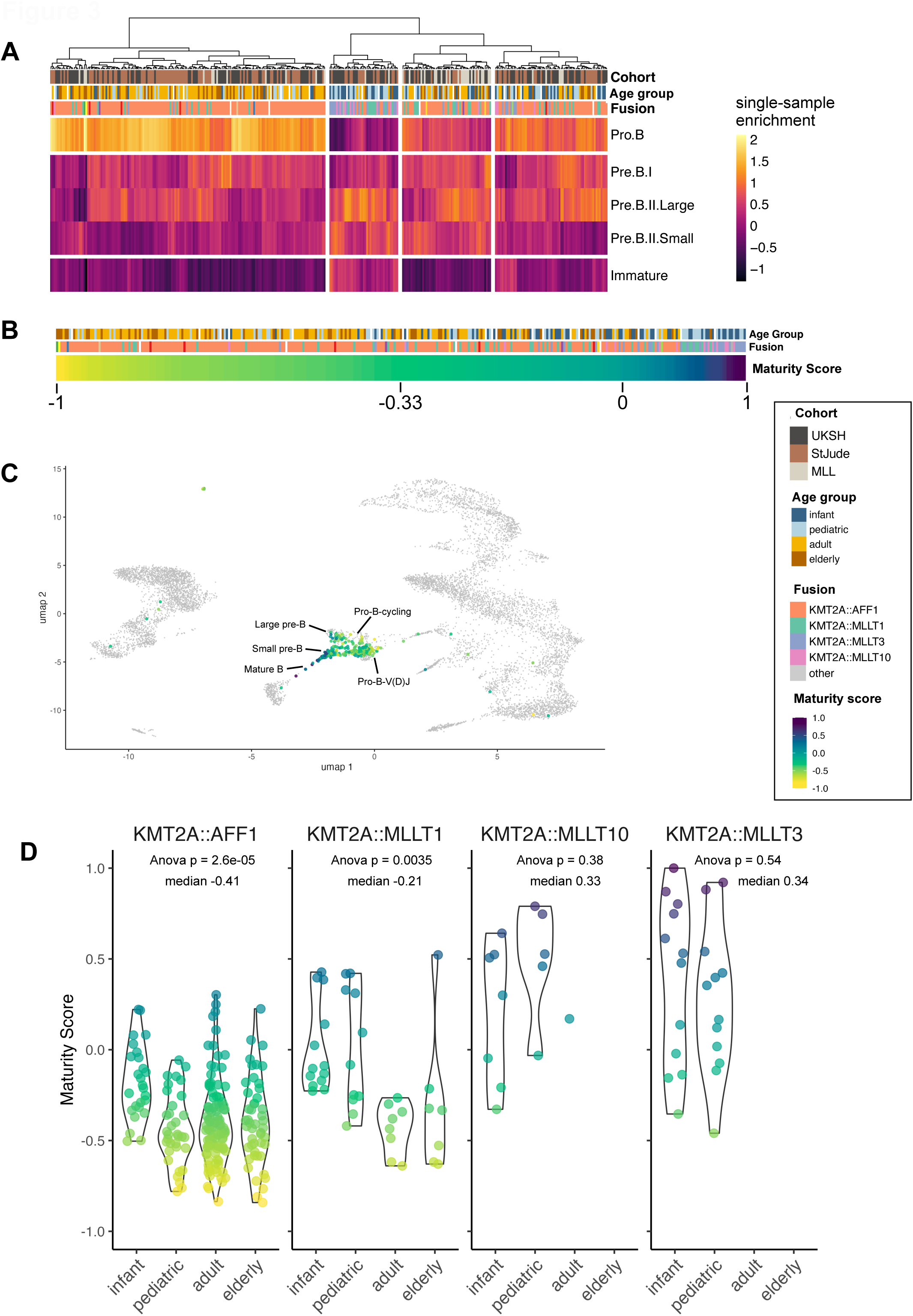
Maturity in KMT2Ar ALL correlates with age and fusion partner. **A**: Gene set enrichment analysis of n=325 *KMT2A*r ALL cases. Enrichment score represents proximity to five healthy B-cell precursor developmental stages (pro-B, pre-B-I, pre-B-II-Large, pre-B-II-Small, Immature-B), calculated based on the RNAseq data using ALLCatchR tool(23). **B**: Developmental trajectories of *KMT2A*r ALL were condensed into a single maturity score ranging from -1 to 1 by fitting linear regression to the enrichments scores in A and calculating the negative slope. Low maturity scores represent immature transcriptional developmental state and proximity to Pro-B cells, high maturity scores represent more mature (pre-B-II, Immature-B) transcriptional developmental states. **C**. *KMT2A*r samples were mapped to the expression matrix of the human single-cell B-cell developmental atlas(24) and colored by maturity score (right). Samples with low maturity score map to early cell stages (pro-B-V(D)J / pre-pro-B cycling), samples with higher maturity score map to mature B-cell stages (small Pre-B / Immature-B). **D**: Maturity score distribution between age groups within driver fusion groups (*AFF1*r n=221, *MLLT1*r n=43, *MLLT10*r n=14, *MLLT3*r n=28). Multi-comparison ANOVA between age groups within each fusion group reveals significantly different maturity scores between age groups in *AFF1*r (p=2.6e-05) and *MLLT1*r (p=0.0035) cases.

We validated the stage of differentiation by correlating *IGH* rearrangement patterns on a genomic level(9) in n=47 patients (n=14 infant, n=13 pediatric, n=20 adult): most patients displayed complete *IGH*-rearrangements, indicative of late pro-B to pre-B cells of origin. In line with previous findings(9), the majority of cases revealed clonally evolving IGH-stems, and the absence of clonal evolution correlated with higher maturity scores (Supplementary Figure S4B). In addition, we mapped bulk-RNAseq data to two single cell atlases of human B-cell development(10,24) confirming the prediction that samples with low maturity scores corresponded to either the pro-B V(D)J compartment or pro-B cycling cells, whereas samples with high maturity scores aligned with more mature B-cell stages. (large pre-B, small pre-B, immature B, Figure 3C and Supplementary Figure S5A). We calculated the “multipotency score” postulated by Iacobucci et al.(10) for our cohort and found a strong correlation with our maturity score (R^2^=0.54, P<0.001, Supplementary Figure 5B) and accordingly, an increase in the maturity score and a decrease in the multipotency score with predicted B-developmental stages based on the B-cell developmental map (Supplementary Figures S5C and S5D respectively).

Increasing maturity correlated with younger age (spearman rank correlation, R=0.45, p<2.2e-15). To separate the effect of age on maturity from the effect of fusion gene, we assessed the maturity scores separately for every driver fusion (Figure 3D). *AFF1*r cases were most immature (n=221, median maturity score=-0.41) compared to samples carrying other fusions. Within *AFF1*r cases, we observed increasing immaturity with increasing age (R=0.16, p=0.024). *MLLT1*r cases had a higher median maturity score than *AFF1*r cases (n=43, median maturity score=-0.21; p=5.3e-06), but again increasing age correlated with increasing immaturity (R=0.6, p=4.8e-05). *MLLT3*r and *MLLT10*r had the highest maturity scores (*MLLT3*r: n=28, median maturity score=0.33; *MLLT10*r: n=14, median maturity score=0.34) and were only observed in young patients (median age 0.95 years and 0.9 years, respectively).

### Age, driver fusion and maturity modulate MRD response

To define the molecular underpinnings of treatment response, we made use of our integrated age-overring cohort including infants (14.5%), pediatric (22.9%), adult (52.8%) and elderly (9.8%) patients treated according to BFM-based pediatric or adult treatment protocols with corresponding MRD measurements available. MRD was measured at two timepoints during induction therapy and after induction/before consolidation using clonal IG/TR or *KMT2A* rearrangements as previously described(25). Of the 214 MRD evaluable patients, 18.2%, 24.8%, and 57.0% had a fast, intermediate, and slow MRD clearance, respectively. Age significantly correlated with the initial MRD clearance (n=214, multi-comparison ANOVA p=0.0002), with the median age of patients in fast MRD responders being 9.6 years in contrast to 37 years in slow responders (Figure 4A). Overall, 51% of infant (n=16/31), 31% of pediatric (n=15/49), 65% of adult (n=73/113) and 86% of elderly (n=18/21) patients showed slow MRD clearance.

**Fig. 4:**
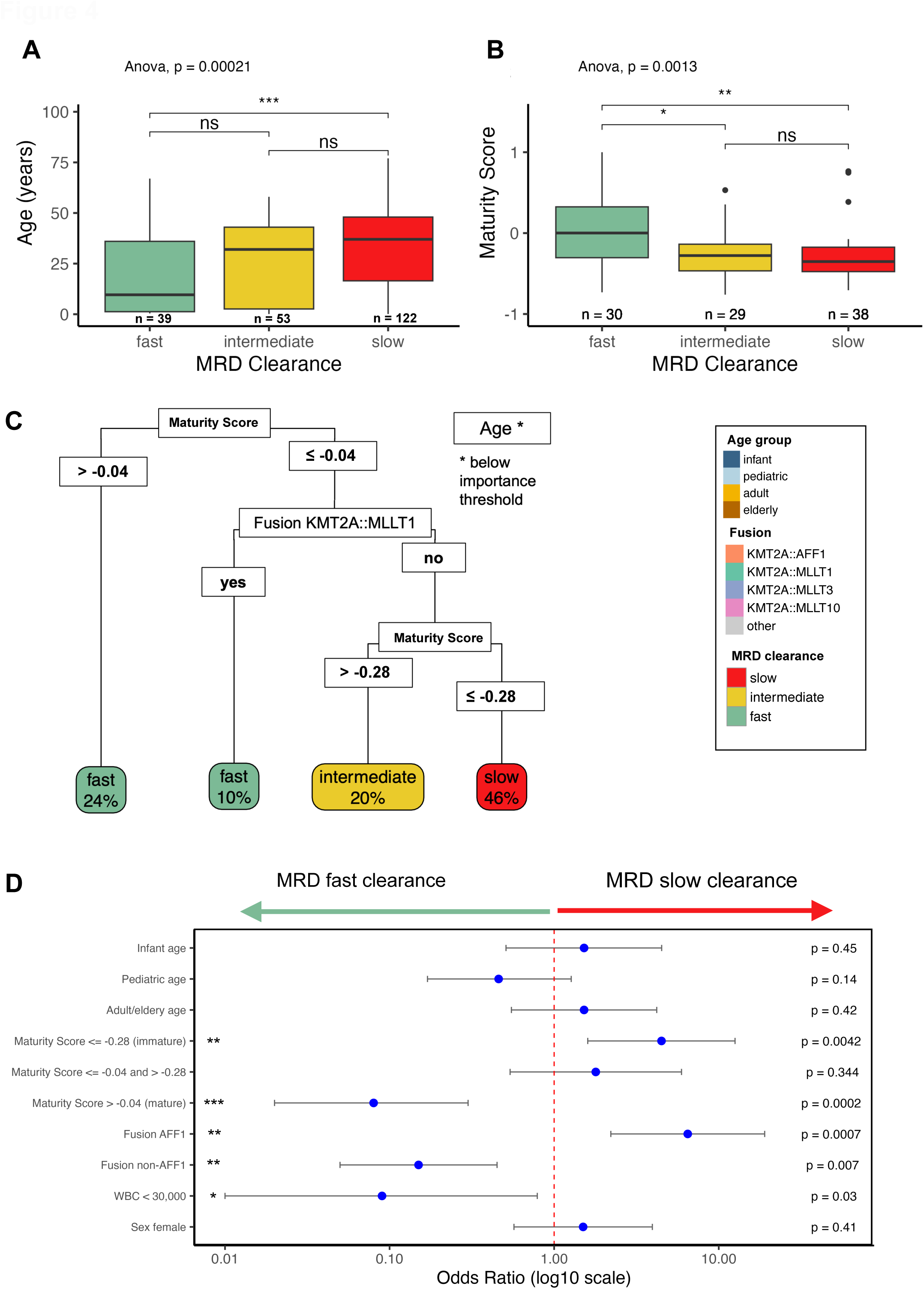
MRD response is modulated by age, gene fusion and maturity score. **A**: Distribution of age within MRD clearance categories (MRD fast clearance: n=39, median age: 9.4 years; MRD intermediate clearance: n=53, median age: 32 years; MRD slow clearance: n=122, median age: 37 years). Age differs significantly higher in MRD slow clearance patients (multi-comparison anova, p=0.00021). **B**: Distribution of maturity scores within MRD clearance categories (MRD fast clearance: n=30; MRD intermediate clearance: n=29; MRD slow clearance: n=398). Maturity scores are significantly higher in MRD fast clearance patients (multi-comparison ANOVA, p=0.0013). **C**: Machine-learning trained decision tree, trained to predict MRD clearance categories based on age, fusion and maturity score. The highest-level split predicts patients with maturity score >-0.04 to have fast clearance, the second split predicts patients with maturity score <=-0.04 and *MLLT1*r to have fast MRD clearance, the third split predicts patients with maturity scores between >-0.28 and -0.04 into intermediate MRD clearance and patients with maturity scores <=-0.28 and no *MLLT1*r to have slow MRD clearance. **D** Univariate analysis of factors predicting MRD clearance (n=97). Categorical factors with multiple levels (age group infant, pediatric, adult/elderly and maturity score group <=-0.28 (immature), <=-0.04 and >-0.28, >-0.04 (mature) were tested per level against all others.

Considering driver fusion partner and maturity as factors correlated with age and each other, we next evaluated MRD clearance with respect to age within each driver fusion group (Figure S6A) and with respect to maturity score (Figure S6B). Considering the most frequent fusion partners *AFF1* (n=144), *MLLT1* (n=23), and *MLLT3* (n=18), we observed a skewed distribution in the three MRD categories fast/intermediate/slow. *AFF1*r cases predominantly showed an intermediate or slow MRD clearance (92%) compared to *MLLT1* (57%) and *MLLT3* (50%) cases with a faster MRD response (total n=185, Chi-squared test with Monte Carlo Simulation, p=0.0005; Supplementary Figure S6A). Correspondingly, infants showed faster MRD response if they were *MLLT1*r or *MLLT3*r, but slower in case of *AFF1*r (*MLLT1*r*/MLLT3*r: 6/13 slow/intermediate; *AFF1*: 17/17 slow/intermediate, Fishers exact test, p=0.0008).

In addition, cases with slow or intermediate MRD clearance had significantly lower maturity scores than patients with fast MRD clearance (Wilcox-test, p=0.0025, Figure 4B). This pattern persisted when considering maturity score, age group and fusion partner separately (Supplementary Figure S6B).

To quantify the respective effects of these factors in a combined analysis, we trained a decision tree (Figure 4C, Material & Methods). The tree grouped patients first by maturity score (>-0.04 to fast), next by fusion (*MLLT1*r to fast) and the remaining cases by maturity score (≤-0.28 to slow, >-0.28 to intermediate), suggesting that the combination of maturity and driver fusion have a high predictive value for MRD response and should be considered together with patient age. We confirmed the impact of maturity, driver fusion and age on MRD clearance in a univariate analysis considering age, sex, fusion partner, maturity score and WBC (Figure 4D).

### MRD driven gene expression signatures

To characterize the molecular programs underlying MRD response, we used RNAseq expression data and built an ordinal regression model, with age and gene fusion as covariates for MRD-associated genes independent of these factors. This resulted in a 448-gene signature (Supplementary Table S6) which grouped patients into three main clusters in an unsupervised clustering approach. Two clusters distinguished patients with fast and slow MRD clearance (Cluster 1; n=15/23 (65% fast clearance, Cluster 3; n=26/27 (96%) intermediate/slow clearance, Fisher’s exact test p=0.007; Figure 5A). We condensed the gene signature into a single “resistance score” using ssGSEA and confirmed that this score was highly predictive for MRD response (Figure 5B).

**Fig. 5:**
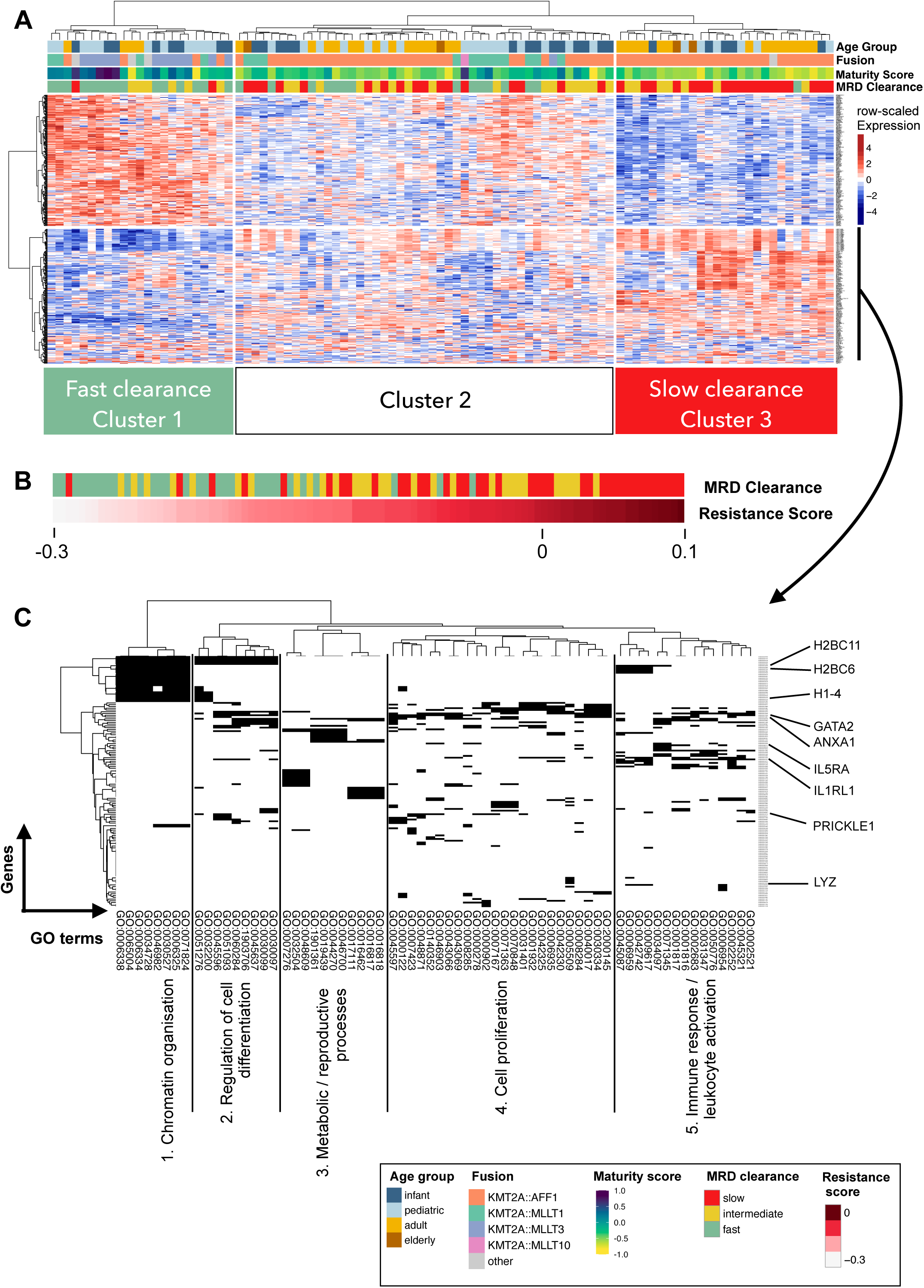
MRD-driven gene expression signatures. **A**: Gene expression signature detected from proportional odds linear regression model grouping patients by MRD response. Cluster 1 is enriched for patients with fast MRD clearance [(Cluster 1: n=12/18 (67%) fast, n=5/18 (28%) intermediate, n=1/18 (6%) slow; Cluster 3: n=1/28(4%) fast, n=7/28 (25%) intermediate, n=20/28 (71%) slow. Chi-squared P<0.0001]). **B:** Resistance score (total score from single sample gene set enrichment analysis using genes upregulated in Cluster 3 as up-set and genes upregulated in Cluster 1 as down-set). **C**: Gene ontology annotation for genes upregulated in MRD Cluster 3. Gene set was annotated with enriched GO terms (GO:Biological Process). Next, enriched GO-terms were filtered for terms size <1000, terms including at least five genes from the gene list and genes were filtered for genes present in at least ten GO-terms. The resulting genes and terms were grouped by functional similarity and five groups were summarized based on included GO-terms.

There was only a minimal overlap of this MRD-gene expression signature with gene expression profiles associated with *KMT2A* fusion partner or defining different B-cell stages (Supplementary Figure S6C) underscoring that the MRD-signature represented specific MRD-associated molecular programs irrespective of the fusion signature.

To functionally annotate the signature of MRD-slow-responders, we applied gene set enrichment analysis for genes upregulated in MRD-slow-responders (Figure 5C) and found enrichment of GO-terms associated with chromatin organization, driven by genes coding for core histones [H1 linker histones *H1-3* and *H1-4*; H2B histones (e.g. *H2BC6, H2BC11*); H2A histones (e.g. *H2AC16*), and H3 histones]. Group 2 and 4 were functionality associated with cell differentiation and proliferation, respectively. The fifth GO-term group was associated with immune response and leukocyte activation and accordingly was driven by immune-modulator genes like *IL5RA, IL1RL1* and *AZU1* and inflammatory response genes like *ANXA1*. Overall, the MRD-associated gene regulation appears not be driven by one prominent signaling pathway but rather reflects a complex interplay of cellular functions related to proliferation and differentiation, which in turn points to the developmental underpinnings (maturity) as a modulator of treatment sensitivity.

### Age, maturity and gene fusion reflect *ex-vivo* drug response

To functionally characterize the MRD response we performed *ex-vivo* DRP on diagnostic primary patient material. We screened 61 *KMT2A*r cases (n=21 infant, n=22 pediatric, n=16 adult, n=1 elderly) with a library of n=27 drugs, including both standard-of-care chemotherapeutics and targeted therapy agents such as BCL2-family inhibitors Venetoclax and Navitoclax, proteasome inhibitor Bortezomib or HDAC-inhibitor Panobinostat (Figure 6A).

**Fig 6.**
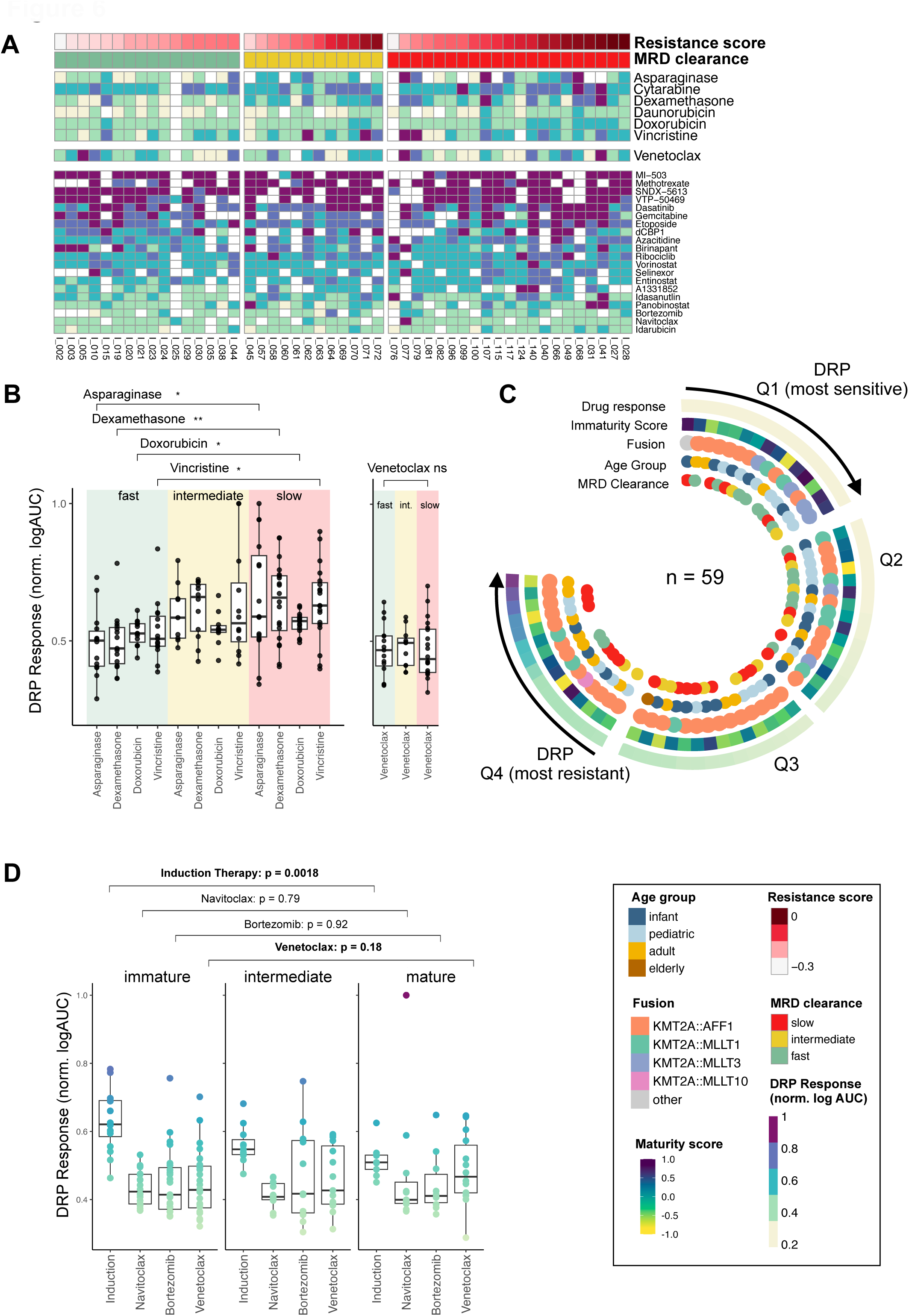
*Ex-vivo* drug response to induction therapy drug mirrors slow MRD clearance. Drug response of *ex-vivo KMT2A*r samples (n=61). **A**: normalized log2 area under the curve (norm.logAUC) for 27 compounds with at least n=40 cases measured(18,19). Top 6 rows represent induction therapy drugs. Patients were grouped according to MRD clearance (n=49) and annotated with resistance score from Figure 5B. **B**: Response (norm.logAUC) to induction therapy drugs was poorer in patients with slow MRD clearance (n=17; red box) compared to patients with fast MRD clearance (n=16; green box; mean norm. log AUC across drugs: 0.61 (slow MRD) vs 0.52 (fast MRD), Wilcox-test per drug: Asparaginase P=0.02, Dexamethasone P=0.0027, Doxorubicin P=0.041, Vincristine P=0.012, Cytarabine and Daunorubicin not significant). **C**: Circos plot of patients (n=59) ordered and separated into quartiles by their mean response (norm.logAUC) to induction phase drugs, and annotated with maturity score, fusion partner, age group and MRD clearance. Quartiles 1 and 2 (Q1/Q2, n=15 respectively; most sensitive) were enriched for fast MRD clearance, high maturity score, non-*AFF1*r and pediatric age. Quartiles 3 and 4 (Q3/Q4, n=15 respectively; most resistant) were enriched for slow MRD clearance, low maturity score, *AFF1*r and older age (Chi-squared tests: MRD clearance P=0.003, maturity score group P=0.064, fusion P=0.153, age group P=0.025, Supplementary Figure S7). **D**: Response to induction phase drugs (left) and Venetoclax (right). Patients grouped by maturity score group (cutoffs -0.04 and -0.28 based on cutoffs from trained decision tree (Figure 4C). Patients in the lowest maturity score group (<=-0.28) are significantly more resistant to patients in the highest maturity Score group (>-0.04) (Wilcox-test, p=0.0018). The effect is inverse for Venetoclax (Wilcox-test p=0.18).

We split samples into quartiles based on their mean drug response to induction phase drugs (Asparaginase, Cytarabine, Dexamethasone, Daunorubicin, Doxorubicin, Vincristine) quantified by the logarithmic area under the dose response curve (logAUC). We observed high concordance with MRD response: samples with a good *ex-vivo* drug response (Q1 and Q2) were significantly enriched for patients with fast MRD clearance (Q1/2 sensitive DRP group included 13/26 fast MRD clearance patients compared to the Q3/4 DRP resistant group with only 2/22 fast MRD clearance patients; Fisher’s exact test p=0.004; Figure 6A, Supplementary Figure S7). This difference in response was also significant for the single drugs Dexamethasone, Daunorubicin, Asparaginase and Vincristine individually (Wilcox-test slow vs. fast MRD clearance; p-value cut-off 0.1; Dexamethasone p=0.0027; Daunorubicin p=0.084, Vincristine p=0.012, Asparaginase p=0.02, Figure 6B).

Q1 and Q2 (most DRP sensitive) were slightly enriched for infant and pediatric patients, although not significantly (Fisher’s exact test p=0.16), significantly enriched for non-*AFF1* gene fusion partners (Fisher’s exact test p=0.01) and had significantly higher maturity scores (Wilcox-test, p=0.016; Figure 6C and Supplementary Figure S7).

In line with these results, our gene expression-based resistance score (Figure 5B), reflecting high expression of MRD-slow-clearance associated genes, was significantly higher in cases with poor response to induction phase drugs (Q1/2: median resistance score -0.13, Q3/4: median resistance score -0.03, Wilcox-test p=0.002). Thus, the activation of MRD-associated gene regulation at diagnosis can be captured by gene expression and is reflected in functional *ex-vivo* drug response profiling (Figure 6A).

We also evaluated three Menin inhibitors (MI-503, SNDX-5613, and VTP-50469). In contrast to our main DRP panel, these compounds were assessed after six rather than three days of incubation to account for their slower kinetics. Within the KMT2A-r cohort, we observed no significant variation in Menin inhibitor sensitivity with respect to developmental maturity, age, or MRD clearance (Supplementary FigureLJS8). For MI-503, AFF1-rearranged cases showed a slightly reduced response compared to other fusions (ANOVA *P*LJ=LJ0.02), although the effect size was modest and not observed for SNDX-5613 or VTP-50469. Because the assay used a different incubation time, these data could not be integrated with the rest of the DRP panel, and we therefore cannot compare overall Menin inhibitor response to other drugs or to different molecular subtypes.

### Venetoclax responders are more frequently immature and characterized by AFF1 fusions

In addition to standard induction chemotherapy drugs, we further analyzed targeted compounds revealing several drugs with notable activities in our cohort including the proteasome-inhibitor Bortezomib (n=45), the HDAC-inhibitor Panobinostat (n=57) and the BCL2-inhibitors Navitoclax (n=46) and Venetoclax (n=58). We matched the sensitivity profiles for these targeted compounds with MRD clearance and maturity to find potential vulnerabilities for MRD-slow responding patients.

By comparing drug response with maturity scores [based on the cutoffs defined in the decision tree in Figure 4C: maturity score <=-0.28 (immature); <=-0.04 and >-0.28, >-0.04 (mature)], we did not observe decreasing sensitivity to Navitoclax, Bortezomib or Venetoclax with increasing immaturity, as opposed to induction therapy drugs (Figure 6D). Interestingly, for Venetoclax we noticed a slight inverse trend showing highest sensitivity in the most immature group, although this trend was not significant (Wilcox-test, p=0.18). Consistently, we did not observe a correlation of MRD clearance with Venetoclax response (Figure 6B). In fact, cases sensitive to Venetoclax mirror the MRD-slow-response profile, with low maturity scores, predominantly *AFF1*r and older age (Supplementary Figure S7), highlighting the potential of Venetoclax in the treatment of cases with MRD-slow response profile, especially those with low B-cell maturity.

## Discussion

Prognosis of *KMT2A*r ALL varies dramatically across age groups, suggesting an intra-subtype molecular heterogeneity to account for the differences in treatment response. To address the underlying biology, we have aggregated the - to our knowledge - largest cohort of *KMT2A*r patients extensively characterized by multi-level data analysis and clinical annotations. This is the first study to integrate transcriptomic developmental state, driver fusion partner, longitudinal MRD response, and *ex-vivo* drug profiling across the full age spectrum of *KMT2A*r B-ALL. By linking these molecular layers to treatment outcome, we identify cellular vulnerabilities and provide a framework for risk stratification. Previous studies(10,7,11–14) on *KMT2A*r ALL have mostly focused on single aspects like age(13), infant patients(7,11,12), or included only small cohorts of *KMT2A*r patients(11,14). Because *KMT2A* fusion partners are unevenly distributed across age groups and influence molecular heterogeneity and outcome, large data sets are necessary to account for these contributing factors(1,2,6). In addition, as only few cooperating genomic events characterize the *KMT2A*r subgroup, it is likely that cell intrinsic factors drive its intra-subtype heterogeneity.

Dissecting the gene expression signatures of *KMT2A*r B-ALLs with respect to fusion partner revealed distinct cellular programs activated by the different driver fusions, indicating a highly proliferative, cell-cycle active gene expression program in *AFF1*r cases and a more B-cell functional, mature signature in non-*AFF1*r cases. Using our newly developed maturity score we observed that older patients exhibited more transcriptionally immature developmental states compared to younger patients, even when controlling for fusion partner. By analyzing maturity within each fusion group and age group, we show that these patterns are consistent across the full age range, suggesting that chronological age and leukemic B-cell maturity are inversely correlated. Age specific difference in *KMT2A*r ALL therefore most likely reflect individual characteristics of a continuous disease spectrum rather than defined sub-entities.

Consistent with prior reports(1,2,4–6), age correlated with early MRD clearance. However, our fine-scale dissection demonstrates that fusion partner and maturity correlate with age and thus a complex interplay of different factors predicts MRD response. Patients with slow or intermediate MRD clearance had significantly lower maturity scores than fast MRD responders, and this association held when maturity, age, and fusion were modeled independently. Integrating these factors as covariates, we derived a novel gene expression signature separating patients by MRD response. Patients with slow MRD clearance displayed activation of cellular programs at diagnosis involved in chromatin organisation, proliferation and immune modulators. Recent studies have begun to link transcriptional maturity to therapy sensitivity in B-ALL. Yoshimura et al.(13) detected a gene-expression based “adult-like” phenotype in pediatric *KMT2A*r, *DUX4*r and *CRLF2*r patients. This phenotype, defined by molecular “adult-likeness” rather than chronological age was linked to decreased Mercaptopurine-sensitivity. Huang et al(14) analysed 203 *ex-vivo* DRPs across molecular subtypes and linked Asparaginase-resistance to pre-pro-B cell of origin, characterized by *BCL2* expression, and suggest Asparaginase and Venetoclax to exhibit synergistic cytotoxicity. Building on these findings, our study expands the analysis to the largest *KMT2A*r cohort, integrating fusion-specific effects, full age range, and direct MRD and *ex-vivo* response correlations.

Functional *ex-vivo* drug profiling mirrored clinical MRD response for drugs administered during initial therapy protocol in line with previous studies(26,27), further advocating the use of *ex-vivo* induction therapy response for risk stratification(28). Integrating our drug response profiling data with the transcriptional profiling revealed that poor response to standard-of-care chemotherapeutics is associated with immature transcriptional states. Accordingly, the MRD-slow-response cases were characterized by an immature developmental stage, *AFF1*r and poorer *ex-vivo* response to standard of care chemotherapeutics. Intriguingly, patients that were responding best to Venetoclax matched the profile of MRD-poor response, with immature developmental stage and predominantly *AFF1*r. Thus, the efficacy of targeted therapies such as Venetoclax was independent of the B-cell developmental states in our functional DRP, suggesting these compounds as a therapeutic alternative in patients with increased risk of poor chemotherapy response.

Using single-cell RNAseq of 89 B-ALL cases (including only nine with *KMT2A*r), Iacobucci et al.(10) developed a “multipotency score”, characterizing patients with an enrichment of early lymphoid progenitors, variation in transcriptional programs and proximity to early B-cell developmental states. The *KMT2A*r cases grouped into a *KMT2A*-early and *KMT2A*-commited cluster, however in-depth characterization of the multipotency of these cases with respect to gene fusion and age is missing. We have reproduced this multipotency score across our cohort and could show that it highly correlates with our maturity score. We extend these insights into a bulk-transcriptome context suitable for clinical application. Together, our findings indicate that leukemic B-cell developmental state and fusion partner jointly shape therapy response and drug sensitivity, exceeding the predictive value of chronological age.

We provide the analytical groundwork to explore interdependencies between transcriptional developmental B-cell state, selection of genomic drivers, gene regulation and biological and clinical phenotypes in B-ALL across molecular subtypes. The inherent correlation of MRD clearance, our MRD-related gene expression signature and the corresponding drug response profile show that molecular programs activated already at diagnosis are essential for and can predict MRD-response as well as are reflected in the functional *ex-vivo* DRP in *KMT2A*r ALL. Our findings highlight the importance of the leukemic B-cell developmental states in determining outcome and drug sensitivity of *KMT2A* rearranged ALL. In future, a consensus definition of B-ALL proximity to normal B-lymphopoiesis might be included in novel risk stratification models, putatively extending beyond *KMT2A*r B-ALL.

## Acknowledgements

RNAseq was performed in cooperation with the Competence Center for Genomic Analysis Kiel (CCGA).

Special thanks to Dr. Rolf Koehler, Institute for Humangenetics, Heidelberg for providing parts of the MRD-data in older children and Dr. Claus Meyer, DCAL, Frankfurt for providing the KMT2A-sequence for MRD analysis.

This study was funded in part by the Deutsche Forschungsgemeinschaft (DFG; German Research Foundation) project number 444949889 (KFO 5010 Clinical Research Unit “CATCH ALL” to A.M.H., L.B., M.J.B, P.E., L.L., N.W., S.B., M.M., M.Br., M.S., G.C, and C.D.B.)

## Authorship

### Contribution

A.M.H., L.B., M.J.B., G.C., C.D.B. designed research;

A.M.H., L.B., M.J.B., J.Ha., E.A., P.E., M.K., T.Be., F.S., K.R., N.W., M.M., B.B. established bioinformatic workflows and performed research;

A.M.H., L.B., M.J.B., L.L., F.S., K.R., B.B., G.C., C.D.B analyzed and interpreted data;

A.M.H., L.B., N.G., A.K.B., M.Br., M.S., G.C., C.D.B. supervised the study;

A.M.H. developed statistical models and produced visualizations.

P.E., S.B contributed RNAseq data;

J.A., A.R., J.He., M.S., G.C., contributed pediatric MRD data;

A.-S.S., T.Bu., N.G. and M.Br. contributed adult MRD data;

J.Ha., E.A., F.S., K.R., B.B., J.-P.B. performed and analyzed drug response profiling;

B.B., J.-P.B. supervised drug response profiling;

C.B. contributed DNA capture panel data;

W.W., C.H. provided samples and clinical and diagnostic data

A.M.H., L.B., C.D.B. wrote the manuscript; and all authors revised and approved the final version of the manuscript

## Supplementary

**Figure S1.**
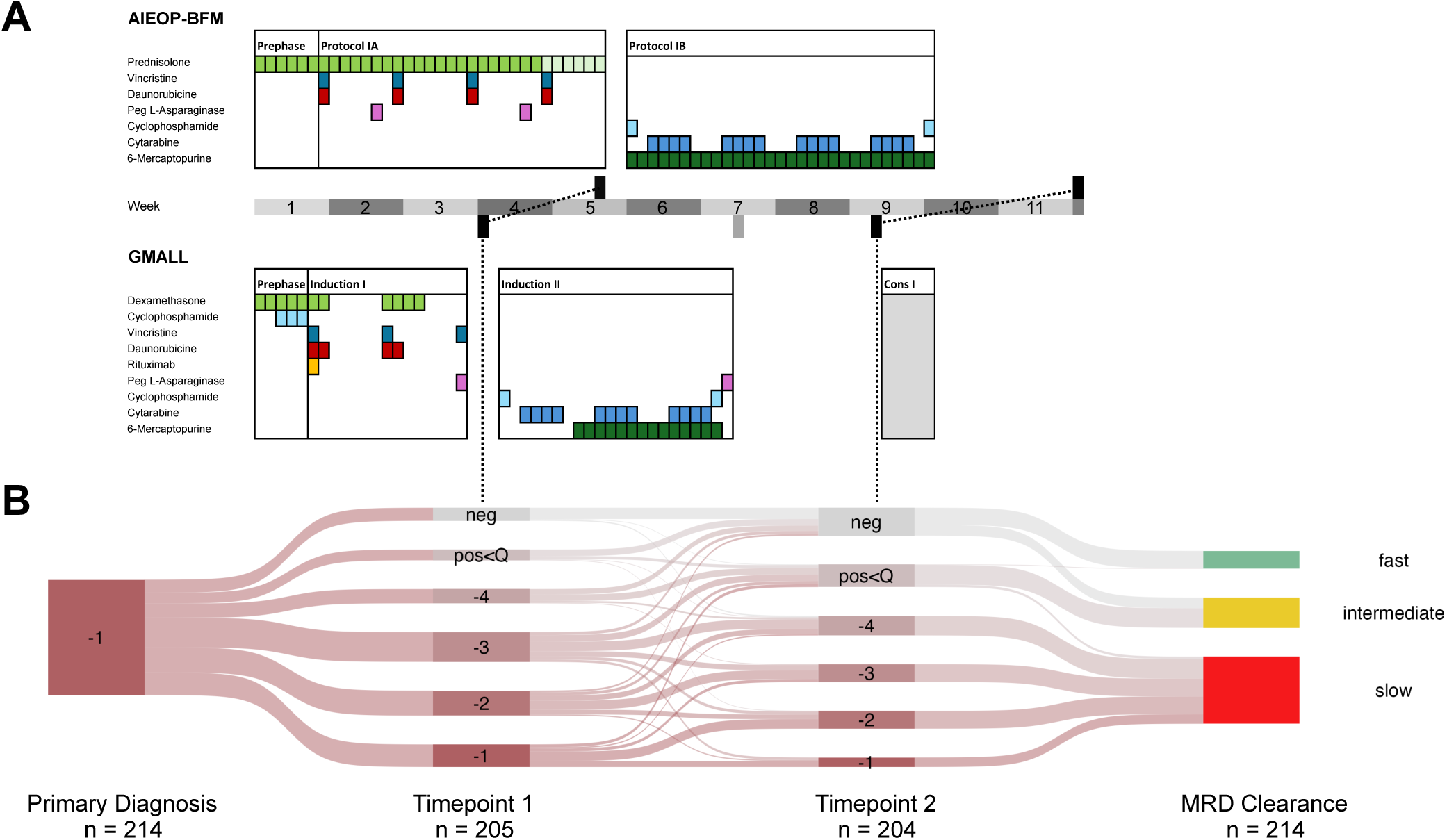

**Figure S2.**
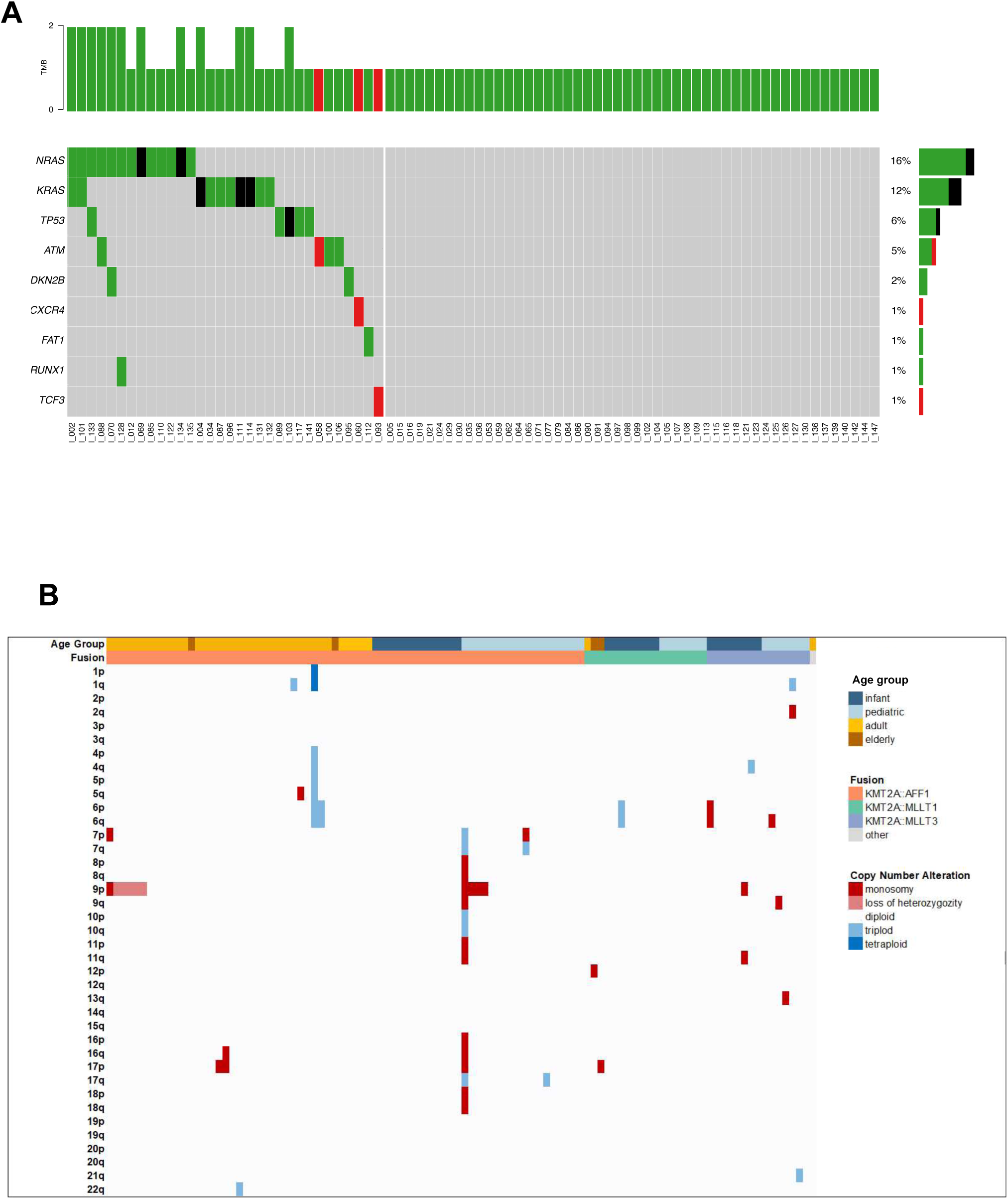

**Figure S3.**
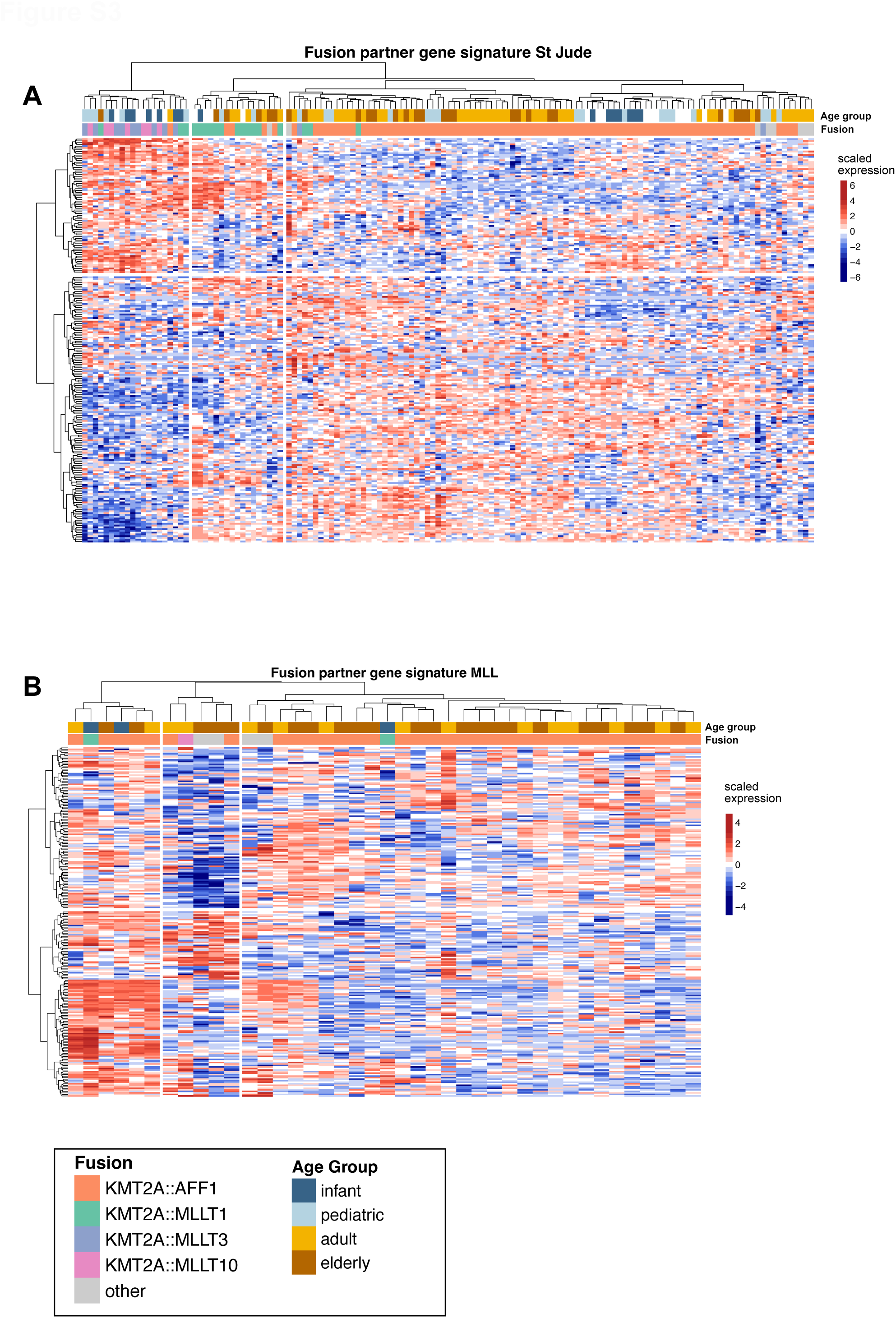

**Figure S4.**
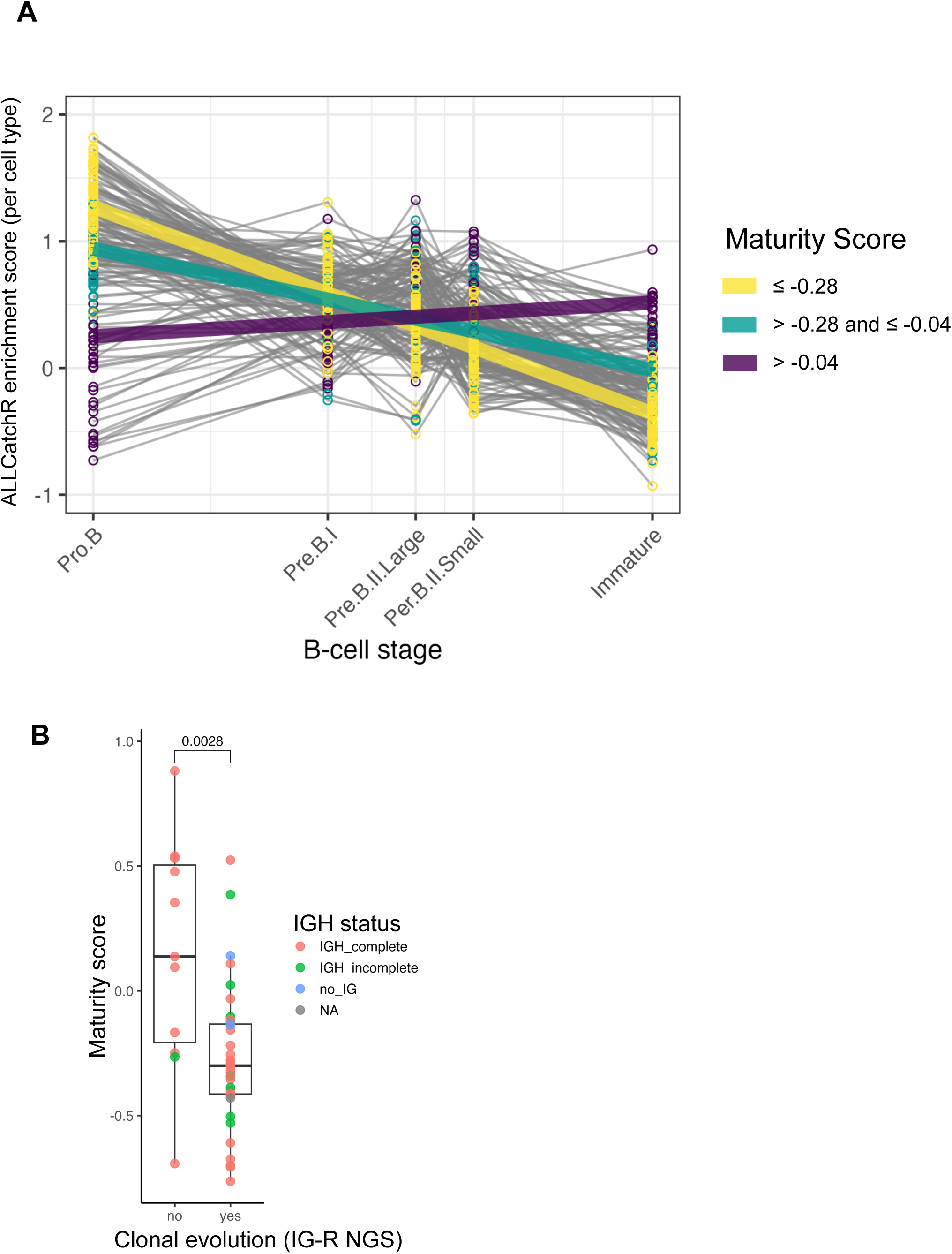

**Figure S5.**
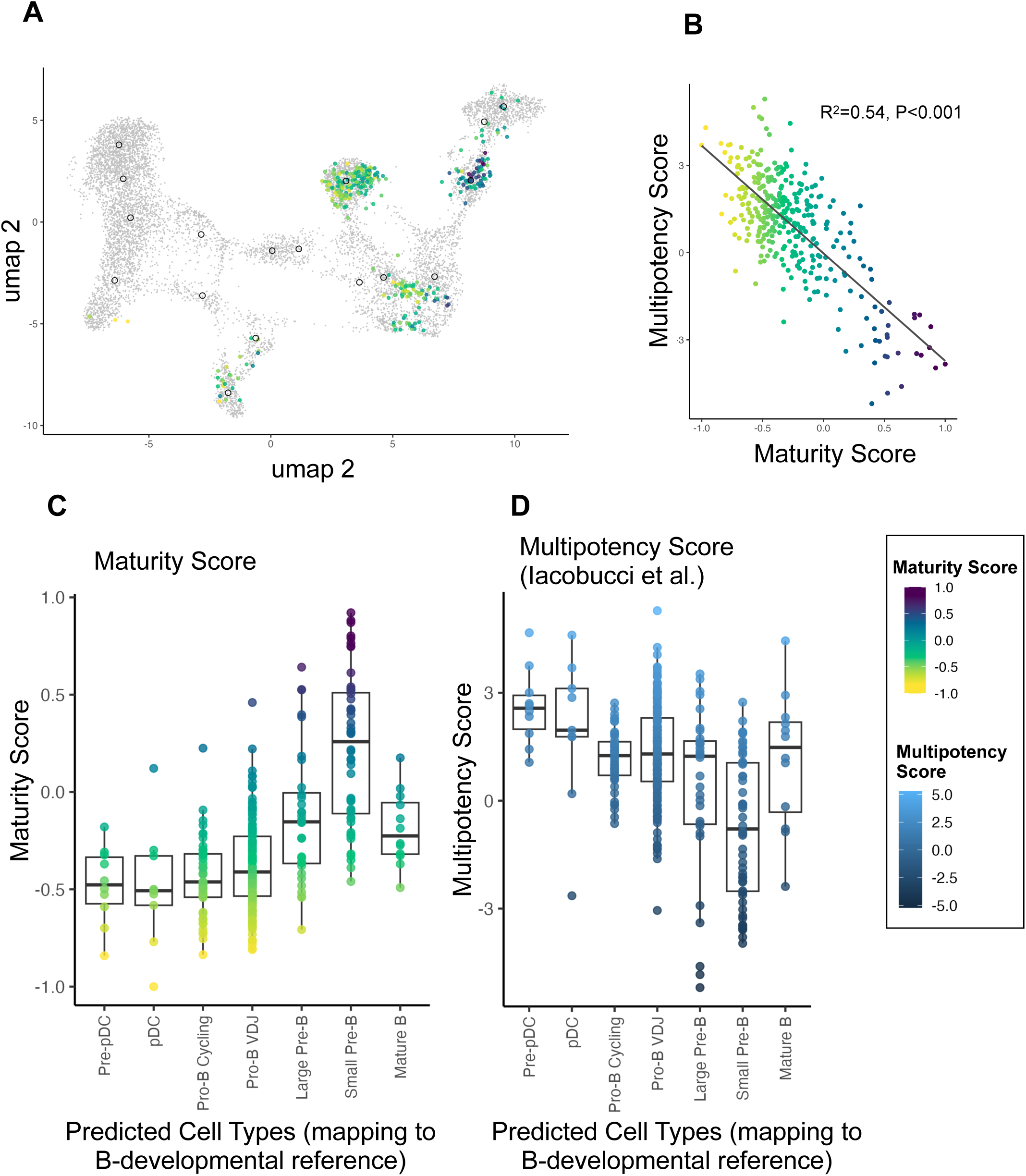

**Figure S6.**
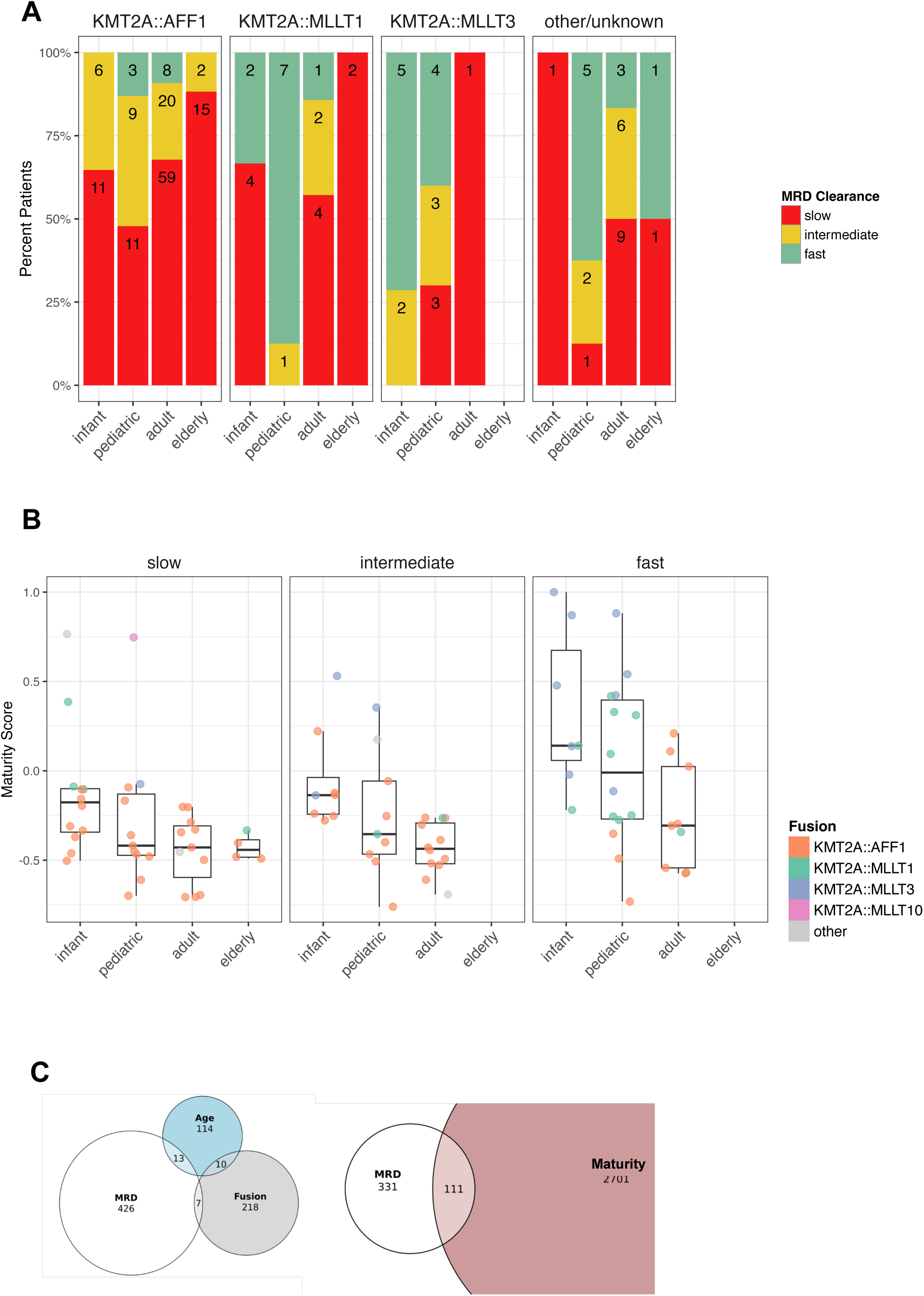

**Figure S7.**
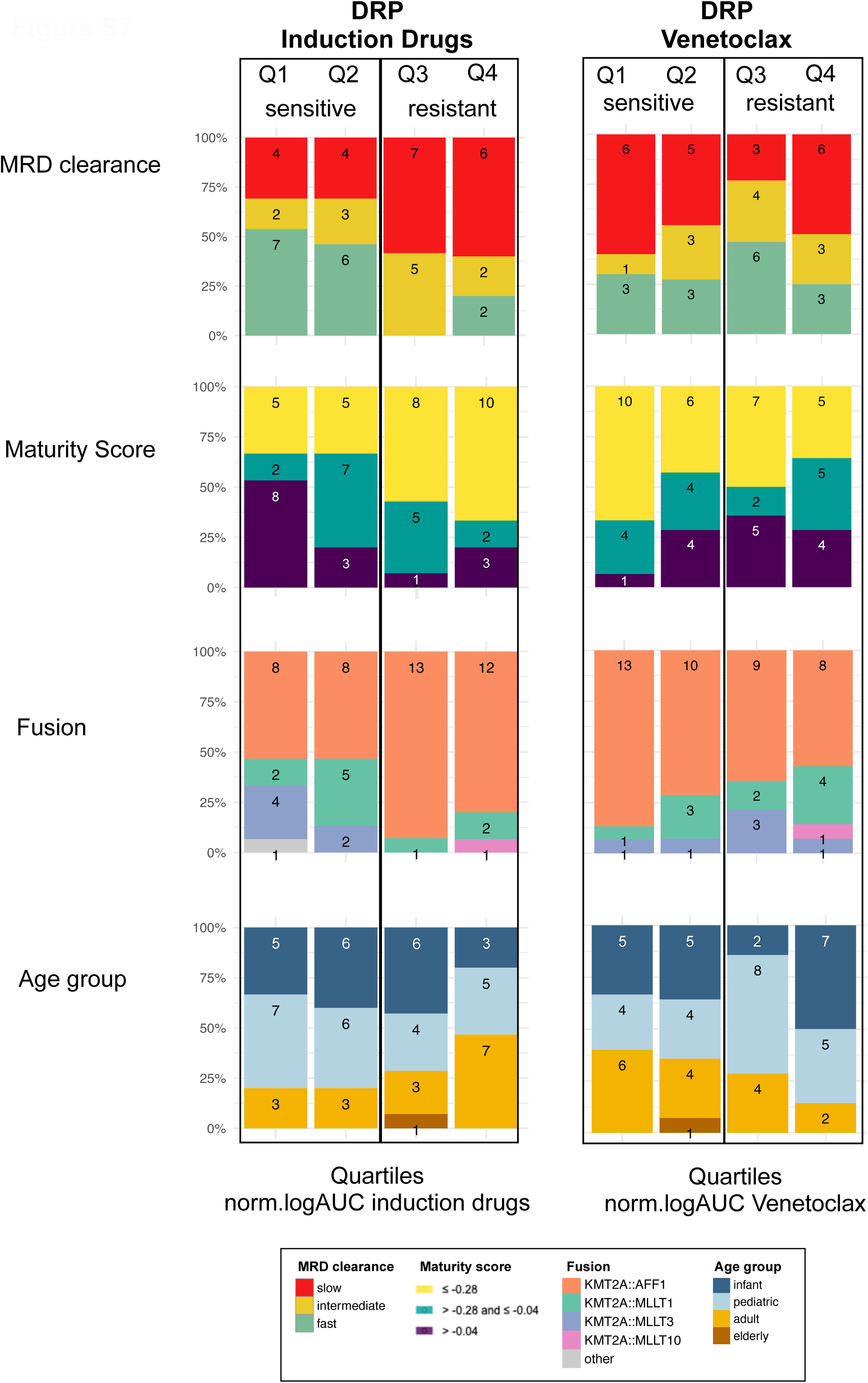

**Figure S8.**
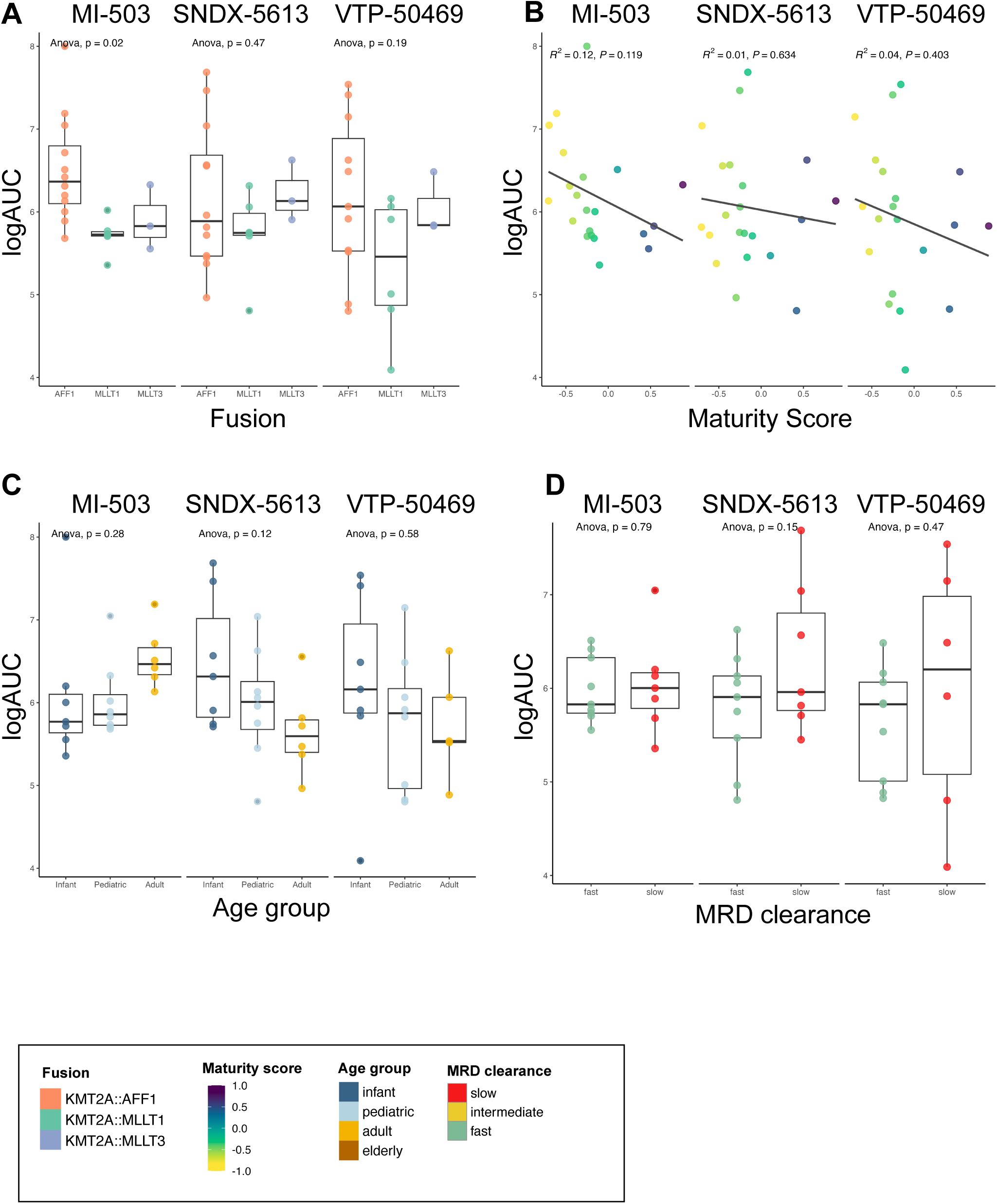

## References

1. Pieters R, Schrappe M, De Lorenzo P, Hann I, De Rossi G, Felice M, et al. A treatment protocol for infants younger than 1 year with acute lymphoblastic leukaemia (Interfant-99): an observational study and a multicentre randomised trial. The Lancet. 2007 July;370(9583):240–50.

2. Szczepański T, Harrison CJ, Van Dongen JJ. Genetic aberrations in paediatric acute leukaemias and implications for management of patients. Lancet Oncol. 2010 Sept;11(9):880–9.

3. Brady SW, Roberts KG, Gu Z, Shi L, Pounds S, Pei D, et al. The genomic landscape of pediatric acute lymphoblastic leukemia. Nat Genet. 2022 Sept;54(9):1376–89.

4. Roberts KG. Genetics and prognosis of ALL in children vs adults. Hematology. 2018 Nov 30;2018(1):137–45.

5. Meyer C, Larghero P, Almeida Lopes B, Burmeister T, Gröger D, Sutton R, et al. The KMT2A recombinome of acute leukemias in 2023. Leukemia. 2023 May;37(5):988–1005.

6. Pui CH, Gaynon PS, Boyett JM, Chessells JM, Baruchel A, Kamps W, et al. Outcome of treatment in childhood acute lymphoblastic leukaemia with rearrangements of the 11q23 chromosomal region. Lancet Lond Engl. 2002 June 1;359(9321):1909–15.

7. Isobe T, Takagi M, Sato-Otsubo A, Nishimura A, Nagae G, Yamagishi C, et al. Multi-omics analysis defines highly refractory RAS burdened immature subgroup of infant acute lymphoblastic leukemia. Nat Commun. 2022 Aug 30;13(1):4501.

8. Kotrova M, Proske C, Darzentas N, Laqua A, Kehden B, Kässens JC, et al. NGS-based IG/TR rearrangement profiling in acute lymphoblastic leukemia: age dependence of immunogenetic maturation. Blood J. 2025 Mar 25;blood.2024027175.

9. Darzentas F, Szczepanowski M, Kotrová M, Kelm M, Hartmann A, Beder T, et al. IGH Rearrangement Evolution in Adult KMT2A-rearranged B-cell Precursor ALL: Implications for Cell-of-origin and MRD Monitoring. HemaSphere. 2023 Jan;7(1):e820.

10. Iacobucci I, Zeng AGX, Gao Q, Garcia-Prat L, Baviskar P, Shah S, et al. Multipotent lineage potential in B cell acute lymphoblastic leukemia is associated with distinct cellular origins and clinical features. Nat Cancer [Internet]. 2025 June 27 [cited 2025 July 25]; Available from: https://www.nature.com/articles/s43018-025-00987-2

11. Khabirova E, Jardine L, Coorens THH, Webb S, Treger TD, Engelbert J, et al. Single-cell transcriptomics reveals a distinct developmental state of KMT2A-rearranged infant B-cell acute lymphoblastic leukemia. Nat Med. 2022 Apr;28(4):743–51.

12. Chen C, Yu W, Alikarami F, Qiu Q, Chen C hui, Flournoy J, et al. Single-cell multiomics reveals increased plasticity, resistant populations, and stem-cell–like blasts in *KMT2A* - rearranged leukemia. Blood. 2022 Apr 7;139(14):2198–211.

13. Yoshimura S, Li Z, Gocho Y, Yang W, Crews KR, Lee SHR, et al. Impact of Age on Pharmacogenomics and Treatment Outcomes of B-Cell Acute Lymphoblastic Leukemia. J Clin Oncol. 2024 Oct 10;42(29):3478–90.

14. Huang X, Li Y, Zhang J, Yan L, Zhao H, Ding L, et al. Single-cell systems pharmacology identifies development-driven drug response and combination therapy in B cell acute lymphoblastic leukemia. Cancer Cell. 2024 Apr;42(4):552–567.e6.

15. Bartram J, Ancliff P, Vora A. How I treat infant acute lymphoblastic leukemia. Blood. 2025 Jan 2;145(1):35–42.

16. Van Der Sluis IM, De Lorenzo P, Kotecha RS, Attarbaschi A, Escherich G, Nysom K, et al. Blinatumomab Added to Chemotherapy in Infant Lymphoblastic Leukemia. N Engl J Med. 2023 Apr 27;388(17):1572–81.

17. Stewart JP, Gazdova J, Darzentas N, Wren D, Proszek P, Fazio G, et al. Validation of the EuroClonality-NGS DNA capture panel as an integrated genomic tool for lymphoproliferative disorders. Blood Adv. 2021 Aug 24;5(16):3188–98.

18. Mihara K, Imai C, Coustan-Smith E, Dome JS, Dominici M, Vanin E, et al. Development and functional characterization of human bone marrow mesenchymal cells immortalized by enforced expression of telomerase. Br J Haematol. 2003 Mar;120(5):846–9.

19. Frismantas V, Dobay MP, Rinaldi A, Tchinda J, Dunn SH, Kunz J, et al. Ex vivo drug response profiling detects recurrent sensitivity patterns in drug-resistant acute lymphoblastic leukemia. Blood. 2017 Mar 16;129(11):e26–37.

20. Gu Z, Churchman ML, Roberts KG, Moore I, Zhou X, Nakitandwe J, et al. PAX5-driven subtypes of B-progenitor acute lymphoblastic leukemia. Nat Genet. 2019 Feb;51(2):296–307.

21. Walter W, Shahswar R, Stengel A, Meggendorfer M, Kern W, Haferlach T, et al. Clinical application of whole transcriptome sequencing for the classification of patients with acute lymphoblastic leukemia. BMC Cancer. 2021 Aug 2;21(1):886.

22. Saorin A, Dehler A, Galvan B, Steffen FD, Ray M, Lu D, et al. Transcriptional remodeling shapes therapeutic vulnerability to necroptosis in acute lymphoblastic leukemia. Blood J. 2025 May 13;blood.2025028938.

23. Beder T, Hansen BT, Hartmann AM, Zimmermann J, Amelunxen E, Wolgast N, et al. The Gene Expression Classifier ALLCatchR Identifies B-cell Precursor ALL Subtypes and Underlying Developmental Trajectories Across Age. HemaSphere. 2023 Sept;7(9):e939.

24. Zeng AGX, Iacobucci I, Shah S, Mitchell A, Wong G, Bansal S, et al. Single-cell Transcriptional Atlas of Human Hematopoiesis Reveals Genetic and Hierarchy-Based Determinants of Aberrant AML Differentiation. Blood Cancer Discov. 2025 Apr 28;OF1– 18.

25. Burmeister T, Ströh AS, Kehden B, Trautmann H, Meyer C, Marschalek R, et al. Measurable residual disease quantification in adult patients with KMT2A-rearranged acute lymphoblastic leukemia. Leukemia. 2024 July;38(7):1600–3.

26. Kaspers GJL, Veerman AJP, Pieters R, Van Zantwijk CH, Smets LA, Van Wering ER, et al. In Vitro Cellular Drug Resistance and Prognosis in Newly Diagnosed Childhood Acute Lymphoblastic Leukemia. Blood. 1997 Oct 1;90(7):2723–9.

27. Enblad AP, Krali O, Gezelius H, Lundmark A, Blom K, Andersson C, et al. Ex Vivo Drug Responses and Molecular Profiles of 597 Pediatric Acute Lymphoblastic Leukemia Patients [Internet]. Oncology; 2024 [cited 2025 Mar 27]. Available from: http://medrxiv.org/lookup/doi/10.1101/2024.12.17.24319138

28. Escherich G, Troger A, Gobel U, Graubner U, Pekrun A, Jorch N, et al. The long-term impact of in vitro drug sensitivity on risk stratification and treatment outcome in acute lymphoblastic leukemia of childhood (CoALL 06-97). Haematologica. 2011 June 1;96(6):854–62.

