## Supplementary figure Legends for "Intra-subtype heterogeneity shapes treatment response in *KMT2A*-rearranged ALL across all age groups"

**Supplementary Figures**

**S1. Treatment schema and MRD clearance classification.** **A**: Induction phase I and II for adult patients (GMALL study group) and pediatric patients (AIEOP-BFM study group) is depicted. **B**: Log levels of MRD measured at the respective reference laboratories using NGS-protocol. All patients had >10% blasts at diagnosis (n=214). Timepoint (TP1) 1 corresponded to “post Induction I” (adult) and post protocol 1A (ped.) respectively (total n=204, of which n=31 infants, n=48 pediatric, n=108 adult and n=18 elderly). Timepoint 2 (TP2) corresponded to “post Induction II”/”pre consolidation I” (adult) and post protocol 1B (ped) respectively (total n=205, of which n=31 infant, n=47 pediatric, n=106 adult, n=20 elderly). Resulting in n=214 patients evaluable for MRD clearance.

**S2.** **Genomic alterations in *KMT2A*r B-ALL**. **A**: Oncoplot showing mutations detected by the lymphoid DNA capture panel in analyzed samples (n=82 patients: n=14 infant, n=12 pediatric, n=50 adult, n=6 elderly). **B**: Karyotypes from SNParrays (n=104) revealed diploid karyotypes in nearly all cases.

**S3. Fusion gene specific gene expression signature applied to validation cohorts.** Heatmap of hierarchical clustering of fusion specific genes detected in supervised analysis using multi-comparison ANOVA and subsequent LASSO feature selection for genes separating *AFF1*r, *MLLT1*r and *MLLT3*r as in Figure 2A. As independent validation, genes were clustered in external cohorts: **A**: St. Jude cohort(1) (n=136) and **B**: Munich Leukemia Laboratory(2) (MLL, n=41).

**S4. Maturity score based on ALLCatchR enrichment scores for proximity to normal B-lymphopoiesis.** To calculate maturity scores per patient, enrichment values for each cell type (y-axis) were plotted against relative cell type distances (x-axis), calculated from group distances in a PCA of immuno-genomic defined RNAseq reference samples of human B-lymphopoiesis(3). A linear regression was fitted through the five data points and the slope of each regression curve was defined as the individual maturity score. **A**: Enrichment values per cell type for all patients (n=325, grey lines) and mean slopes for patients belonging to the three maturity score groups (<=-0.28 (immature): n=186; <=-0.04 and >-0.28: n=74, >-0.04 (mature): n=65) as defined by the trained decision tree (Figure 4C). **B:** Maturity scores with respect to presence of clonal evolution from IG-R NGS sequencing (n=47 samples of which n=9 infant, n=9 pediatric, n=16 adult; Wilcox-test p=0.0028).

**S5. Validation of maturity score by mapping to single-cell B-cell developmental map. A:** Predicted per sample cell types and projection to umap. Each bulk RNA-sample was projected to the B-cell developmental map(4) using the BoneMarrowMap R package(5). Projected samples are colored by maturity score value. **B:** Per sample Maturity Score (x-axis) vs. Multipotency Score as postulated by (4) and correlation of scores (Pearson correlation R^2^=0.54, P<0.001). Solid line represents linear regression curve. **C:** Boxplots of per patient Maturity scores per predicted B-cell developmental map cell type. **D:** Boxplots of per patient Multipotency scores per predicted B-cell developmental map cell type.

**S6. Maturity score correlates with fusion and age groups and differs between MRD clearance groups.** **A**: Fine-scale pattern of MRD clearance categories per age group and fusion including n=144 *AFF1*r, n=23 *MLLT1*r, n=18 *MLLT3*r and n=28 cases with other or unknown fusion. **B**: Distribution of maturity Score (range: -1 to 1) within MRD clearance categories (MRD fast clearance: n=30, MRD intermediate clearance: n=29; MRD slow clearance: n=38, ANOVA: p=0.0013, Wilcox-test between groups: fast vs intermediate p=0.013, fast vs slow p=0.0025, intermediate vs slow p=0.65. **C**: Venn diagram of gene set intersection between MRD score genes, Fusion-specific genes as described in Fig 2A and genes significantly correlated with age and MRD score genes and genes used in the ALLCatchR tool(3) for proximity scores to normal B-cells.

**S7. Enrichment for MRD clearance category, maturity score, fusion and age group per induction phase and Venetoclax response quartile.** Patients were split into quartiles based on mean response to induction phase drugs (Asparaginase, Cytarabine, Daunorubicin, Dexamethasone, Doxorubicin, Vincristine; n=59) or Venetoclax (n=58). Left side: DRP induction phase drugs. MRD clearance (fast vs. intermediate/slow clearance, Q1/2: vs. Q3/4, Fisher’s exact test p=0.004), maturity scores (<=-0.28 (immature); <=-0.04 and >-0.28, >-0.04 (mature), Q1/2 vs. Q3/4 Chi-sq. test p=0.055), fusion partners (*AFF1* vs non-*AFF1* Q1/2: vs. Q3/4, Fisher’s exact test p=0.01), age groups (infant/pediatric vs adult/elderly, Q1/2 vs. Q3/4, Fisher’s exact test p=0.16). Right side: DRP Venetoclax. MRD clearance (fast vs. intermediate/slow, Q1/2 vs. Q3/4, Fisher’s exact test p=0.75), maturity scores (<=-0.28 (immature); <=-0.04 and >-0.28, >-0.04 (mature), Q1/2 vs. Q3/4, Chi-sq. test p=0.41), fusion partners (*AFF1* vs non-*AFF1*, Q1/2 vs. Q3/4, Fisher’s exact test p=0.15) age group (infant/pediatric vs adult/elderly, Q1/2 vs. Q3/4, Fisher’s exact test p=0.25).

**S8.** **Menin-inhibitors do not show significant intra-subtype heterogeneity in *KMT2A*r B-ALL.** LogAUC values for Menin-Inhibitors MI-503, SNDX-5613 and VTP-50469 with respect to **A:** Fusion (ANOVA p values denoted), **B**: Maturity score (pearson correlation coefficient and p-value noted, solid line represents linear regression), **C:** Age group (ANOVA p-values denoted) and **D:** MRD clearance. Boxes with less than 3 data points were removed.

1.	Gu Z, Churchman ML, Roberts KG, Moore I, Zhou X, Nakitandwe J, et al. PAX5-driven subtypes of B-progenitor acute lymphoblastic leukemia. Nat Genet. 2019 Feb;51(2):296–307.

2.	Walter W, Shahswar R, Stengel A, Meggendorfer M, Kern W, Haferlach T, et al. Clinical application of whole transcriptome sequencing for the classification of patients with acute lymphoblastic leukemia. BMC Cancer. 2021 Aug 2;21(1):886.

3.	Beder T, Hansen BT, Hartmann AM, Zimmermann J, Amelunxen E, Wolgast N, et al. The Gene Expression Classifier ALLCatchR Identifies B-cell Precursor ALL Subtypes and Underlying Developmental Trajectories Across Age. HemaSphere. 2023 Sept;7(9):e939.

4.	Iacobucci I, Zeng AGX, Gao Q, Garcia-Prat L, Baviskar P, Shah S, et al. Multipotent lineage potential in B cell acute lymphoblastic leukemia is associated with distinct cellular origins and clinical features. Nat Cancer [Internet]. 2025 June 27 [cited 2025 July 25]; Available from: https://www.nature.com/articles/s43018-025-00987-2

5.	Zeng AGX. _BoneMarrowMap: Single cell reference mapping onto Bone Marrow	Hematopoiesis_. 2024.
