## Supplementary Methods for "Intra-subtype heterogeneity shapes treatment response in *KMT2A*-rearranged ALL across all age groups"

**Library preparation and sequencing of discovery cohort**

RNA was extracted from primary bone marrow aspirates or blood with at least 20% blasts at initial diagnosis and sequenced according to published protocols(1). Gene counts were normalized in R (Version 4.3.0) using the package DGEobj.utils (Version 1.0.6) to normalized log2 counts per million.

**Fusion gene detection**

Gene fusions were detected by PCR (n = 85) or from the transcriptome data using the fusioncatcher tool (Version 1.33).

**DNA capture sequencing**

Single nucleotide variations were called from genomic capture panels. Variants were annotated using variant effect predictor and filtered for pathogenic somatic mutations.

**Clinical Response Data**

We defined two early MRD assessment timepoints at which patients of all age groups had received comparable treatment protocols (Supplementary Figure S1A). Timepoint 1 (TP1) corresponding to “MRD End of Induction” in the pediatric protocol (main components: Glucocorticoids, Daunorubicin, Vincristine, Asparaginase) and to “before Induction II” in the adult treatment protocol. Timepoint 2 (TP2) corresponded to “MRD End of consolidation” (main components: Cyclophosphamide, Cytarabine, 6-Mercaptopurine) in the pediatric protocol and to “post Induction II” (main components: Cyclophosphamide, Cytarabine, 6-Mercaptopurine, Asparaginase) in the adult protocol. Thus, we classified MRD evaluable patients based on MRD kinetics into “Fast MRD clearance” if MRD was negative or positive below quantifiable range at TP1 and negative at TP2. “Slow MRD clearance” was defined as positive MRD (at least -04) at TP2. Patients with positive MRD at TP1 and negative MRD or positive below quantifiable range MRD at TP2 were defined “intermediate” (Supplementary Figure S1B, Supplementary Table S5).

**Drug response profiling**

Leukemia cells were seeded on an hTERT-immortalized mesenchymal stromal monolayer and incubated with a compound library encompassing both FDA-approved, clinically actionable chemotherapeutics as well as single agents targeting specific signaling pathways and molecular targets (ABL/Src, BCL2, proteasome, HDAC, menin). Cells were exposed to the drugs in a six-point serial dilution for 72h before reading out their viability by fluorescence microscopy (Operetta, Perkin Elmer) using CyQuant as a live-cell marker. Viable cells were counted per condition and normalized to DMSO. The drug responses were fitted to a three-parameter dose-response model (logEC_50_, *E*_max_, slope parameter *b*) using the *drc* R package (Ritz et al. PlosOne, 2015) and integrated to yield the area-under-the-curve (AUC) as a drug scoring metric for estimating sensitivity or resistance(2,3).

**MRD gene expression signature**

The MRD gene expression signature was defined using an ordinal regression model to account for the structured nature of MRD categories. The model was calculated in R using the proportional odds logistic regression model in the MASS package (Version 7.3-60.0.1). For each coding gene, we calculated the effect of gene expression on the MRD category including the driver fusion and age as variables in the model. Significance scores were calculated by comparing the model t-values for the effect of gene expression on MRD to the normal distribution and log2-fold-changes which were calculated between MRD categories “fast” and “slow”. Genes were filtered for p-values smaller or equal to 0.05 and a log2-fold-change ≧1 (Supplementary Table S6).

**Gene set enrichment analysis**

MRD signature

To characterize the genes that were associated with slow MRD clearance, we extracted the genes upregulated in the respective gene expression cluster and searched for enrichment in functional GO-terms. We next filtered for genes most abundant in cell functional modules (present in at least 10 GO-terms) and for GO-terms including at least five genes from the list. We grouped genes and the respective enriched GO-terms which lead to five GO-term groups that represent diverse functionalities of the cell.

**Maturity score calculation**

We fitted linear slopes on the enrichment across B-cell developmental stages calculated by the ALLCatchR tool(4).

To define relative distances between B-cell stages we used PCA projections of cell type predictions on healthy B-cell populations from FACS sorted RNAseq samples to calculate the relative distances in a 2D-projection. For each individual patient, we calculated the linear slope fitted to the enrichment scores for the five B-cell stages (Pro-B, Pre-B-I, Pre-B-II-Large, Pre-B-II-Small, Immature-B) vs their relative distances, resulting in a single-value estimate. The slope defines the maturity score. A steep positive slope value indicated low enrichment for early cell stages and high enrichment for more mature cell stages and hence corresponds to high maturity, whereas patients with steep negative slopes had high enrichment for earlier cell types and low enrichment for mature cell stages and thus an overall low maturity (Supplementary Table S3). For validation, we mapped our bulk RNA-seq cohort to two single-cell bone marrow atlases of physiological B-cell development(5,6). We used the BoneMarrowMap R package(7) to predict most likely cell stage per bulk sample from the respective data sets and to map bulk samples to the single cell reference.

Validation by IG-R characterization

Somatic recombination at the IGH-locus can be separated in DH to JH recombination, VH to DJ recombination and VH replacement and these steps happen successively during normal B-cell development. Aberrant ongoing D to J and V to DJ recombination has been described in leukemia and was found in most of our samples. We detected clonal evolution as described previously(8).

**Impact of fusion, immaturity and age on MRD clearance**

Decision Tree

with 10-fold cross validation and a maximum tree depth of five. We tested robustness across five different seeds.

Univariate Analysis

We calculated odds ratios of each age group (infant, pediatric, adult/elderly) compared to all others, gene fusion (*AFF1*r, non-*AFF1*r), white blood cell count over and under 30,000, and immaturity scores as categorical values based on cutoffs found in the decision tree and sex.

1.	Bastian L, Hartmann AM, Beder T, Hänzelmann S, Kässens J, Bultmann M, et al. UBTF::ATXN7L3 gene fusion defines novel B cell precursor ALL subtype with CDX2 expression and need for intensified treatment. Leukemia. 2022 June;36(6):1676–80.

2.	Frismantas V, Dobay MP, Rinaldi A, Tchinda J, Dunn SH, Kunz J, et al. Ex vivo drug response profiling detects recurrent sensitivity patterns in drug-resistant acute lymphoblastic leukemia. Blood. 2017 Mar 16;129(11):e26–37.

3.	Saorin A, Dehler A, Galvan B, Steffen FD, Ray M, Lu D, et al. Transcriptional remodeling shapes therapeutic vulnerability to necroptosis in acute lymphoblastic leukemia. Blood J. 2025 May 13;blood.2025028938.

4.	Beder T, Hansen BT, Hartmann AM, Zimmermann J, Amelunxen E, Wolgast N, et al. The Gene Expression Classifier ALLCatchR Identifies B-cell Precursor ALL Subtypes and Underlying Developmental Trajectories Across Age. HemaSphere. 2023 Sept;7(9):e939.

5.	Zeng AGX, Iacobucci I, Shah S, Mitchell A, Wong G, Bansal S, et al. Single-cell Transcriptional Atlas of Human Hematopoiesis Reveals Genetic and Hierarchy-Based Determinants of Aberrant AML Differentiation. Blood Cancer Discov. 2025 Apr 28;OF1–18.

6.	Iacobucci I, Zeng AGX, Gao Q, Garcia-Prat L, Baviskar P, Shah S, et al. Multipotent lineage potential in B cell acute lymphoblastic leukemia is associated with distinct cellular origins and clinical features. Nat Cancer [Internet]. 2025 June 27 [cited 2025 July 25]; Available from: https://www.nature.com/articles/s43018-025-00987-2

7.	Zeng AGX. _BoneMarrowMap: Single cell reference mapping onto Bone Marrow	Hematopoiesis_. 2024.

8.	Darzentas F, Szczepanowski M, Kotrová M, Kelm M, Hartmann A, Beder T, et al. IGH Rearrangement Evolution in Adult KMT2A-rearranged B-cell Precursor ALL: Implications for Cell-of-origin and MRD Monitoring. HemaSphere. 2023 Jan;7(1):e820.
